# A Two-Fluid Model of Brain Dynamics

**DOI:** 10.64898/2026.06.25.734626

**Authors:** Ahmed Farag Ali, Nader Inan, Ruben Laukkonen, Pavlo Mikheenko

## Abstract

We develop a theoretical proposal linking vacuum stability and brain dynamics through superconductivity-inspired coherence, symmetry reduction, and the thermodynamic stabilization of low-entropy regimes. We take an unbroken SU(3) structure as a candidate stable residue of the low-temperature vacuum. At the neural level, we formulate a coarse-grained analog in which a two-fluid model with dissipative and coherence-supporting components describes brain dynamics. Specifically, the coherence-supporting component is proposed as a possible basis for the efficient binding and integration required to sustain a stable, unified conscious state. The proposal offers a common geometric language for relating physics and neuroscience with falsifiable signatures in coherence and state-dependent transitions. The main technical contribution is a computational algebraic model of conscious-state dynamics, where neural data are mapped to reconstructed state trajectories. Effective generators are inferred from those trajectories, and the two-fluid split is tested as a Cartan-root decomposition of *su*(3), with a rank-two commuting sector for coherence-preserving balance and six root directions for state transitions. This structure can be tested on neural data and contrasted with alternative dynamical models.

## 1 Introduction

The provocative hypothesis that biological systems, especially neural substrates such as the human brain, might support superconductivity at or near physiological temperatures presents significant interdisciplinary implications for physics and neuroscience, with direct relevance to consciousness studies. Recent experiments from a single research programme have reported superconductivity-like signatures in microtubules, the protein filaments densely populating neurons, including Meissner-type magnetic screening, vortex-like magnetic structures, and Josephson-junction-like transport features [1]. Related measurements on excised brain slices have reported abrupt resistance changes, gap-like anomalies, and nonlinear current-voltage characteristics that have been interpreted as possible superconducting behavior, although independent replication is needed [2].

Rigorous models linking physics and consciousness remain in their early stages, but recent work on brain thermodynamics and quantum biology has begun to supply relevant conceptual and empirical tools [3, 4]. For example, whole-brain modeling work has found that human brain dynamics display a quantum-like probability structure, including interference-like effects in coupled-oscillator systems, suggesting that quantum-formal mathematical tools can organize classical neural dynamics without requiring the brain to be a literal quantum computer [5]. This provides an important independent motivation for the present algebraic approach: whether reconstructed neural state spaces possess a precise computational structure. More broadly, the two-fluid model (including superconductivity-like coherence) can, in theory, provide a candidate physical mechanism for several unresolved questions in neuroscience:

1. **Rapid long-range coordination:** how spatially distributed neural populations coordinate their activity rapidly and selectively despite finite conduction delays, biological noise, and continuously changing sensory input. [6]
2. **Binding and conscious unity:** how features processed across different neural populations are dynamically bound into coherent perceptual objects and integrated into a stable, continuously updated model of the self and world. [7]
3. **Energetically efficient computation:** how the brain sustains complex, large-scale information processing within a highly constrained metabolic budget, given the substantial energetic costs of synaptic transmission. [8]
4. **The balance between integration and flexibility:** how the brain maintains globally integrated and relatively stable organization while preserving local specialization, differentiation, and the capacity to reconfigure rapidly as cognitive demands change. [9]

Within the present hypothesis, these challenges are approached through superconductivity-inspired coherence as a candidate model rather than an established biological mechanism. Superconductive coherence enables spatially distributed components to participate in a common collective state, supporting rapid coordination and the binding of differentiated signals into an integrated whole. Its low-dissipation dynamics provide a model for sustaining large-scale computation within tight energetic constraints. Moreover, the coexistence of coherent and dissipative components preserves the flexibility required for local specialization, adaptive reconfiguration, and transitions between cognitive states.

Superconductivity, first observed by Kamerlingh Onnes in 1911 through the vanishing electrical resistance of mercury [10], is a macroscopic quantum state characterized by zero dc resistance and magnetic flux expulsion below a critical temperature, as later established by the Meissner–Ochsenfeld effect [11]. For conventional superconductors, the microscopic Bardeen– Cooper–Schrieffer (BCS) theory explains superconductivity through the formation of electron pairs, known as Cooper pairs, which collectively condense into a macroscopic quantum coherent state, exhibiting dissipationless currents [12]. Modern research into dense hydrogen-rich compounds has shown that strong electron–phonon coupling involving light hydrogen modes can support very high superconducting transition temperatures, but only under extreme pressure in the best-established cases [13, 14]. Notably, biological systems, including the brain, contain abundant hydrogen in water and organic molecules; hence the structured water, confined geometry, and ordered charge environment of the brain may motivate superconductivity-like hypotheses [15].

In neurons, microtubules are ordered nanoscale protein polymers with internal cavities and confined water. These features have motivated proposals for unconventional collective transport, partly inspired by Little’s proposal of high-temperature superconductivity in low-dimensional organic systems [16] and later extensions to superconducting or superconducting-like behavior in microtubules [1, 2]. Such a low-dissipation channel could influence neural synchronization and long-range coordination. Moreover, superconducting neuromorphic systems offer useful technological analogies [17, 18]. Claims of biological superconductivity at or above room temperature clearly require independent replication, including: magnetic and thermodynamic tests, field-dependent transport measurements, and exclusion of ionic conduction, electrode artifacts, sample damage, and non-superconducting coherence effects.

At the symmetry level, superconductivity arises when a charged condensate develops phase rigidity. This rigidity gives the electromagnetic gauge field an effective mass through the Anderson-Higgs mechanism and produces the Meissner effect, in which magnetic fields penetrate only over a finite London length. Zero dc resistance follows from the protected current response of the ordered condensate. In the low-energy framework developed here, the remnant unbroken symmetry is SU(3), and the neural model uses this residual structure to organize the two-fluid picture. It therefore treats SU(3) as an organizing principle for bound, non-factorizing collective dynamics. Major theories of consciousness agree that experience appears as a unified whole. Integrated Information Theory frames this as irreducibility [19], global-workspace theory as system-wide availability [20], and binding-based neuroscience as coordinated integration [21, 22]. In this sense, consciousness is many processes held together as one experience, and SU(3) is introduced here as a compact non-factorizing structure for modeling such unity.

In this work, we propose a layered picture connecting superconductivity, the Free Energy Principle, the two-fluid model, and an organizing geometry associated with SU(3). The two-fluid model serves as a neural analogy: ordinary neural activity forms one component, while binding and integration capacity may arise as a state-dependent resource in the other. We develop a possible SU(3)-inspired neural geometrodynamics by treating SU(3) as a hypothesis about reconstructed neural dynamics. By integrating these ideas, the paper offers a shared geometric language for neuroscience and physics. The layered analogy is introduced after the superconductivity and quasi-one-dimensional context has been established. The principal open empirical question is whether reconstructed neural dynamics can be organized into an *su*(3) structure, with two slow commuting modes and six corresponding transition dynamics. If so, this structure could support a new neural-manifold view of consciousness based on Lie-algebraic reconstruction.

## 2 The Free Energy Principle and Spontaneous Symmetry Breaking of U(1) Gauge Symmetry

The Free Energy Principle (FEP), formulated by Karl Friston, offers a comprehensive framework for understanding how biological systems maintain their internal states and adapt to external environments by minimizing a free energy functional [23]. This principle posits that organisms strive to reduce the discrepancy between their internal models and sensory inputs, thereby achieving a state of sustained equilibrium (i.e., homeostasis and alostasis). To explore the intersection of the FEP with fundamental physics, we examine the Ginzburg-Landau (GL) theory of superconductivity.

However, it is important to clarify that while the GL free energy arises from physical and thermal processes, the FEP concerns a *variational free energy* defined over probability distributions in an informational sense. Foundational holographic and boundary-based frameworks establish deep links between geometry and entropy in gravitational settings [24–26]. Even so, one must exercise caution in directly equating the FEP’s free energy with the thermodynamic free energy of superconductivity; the former is inferential and statistical, whereas the latter is explicitly thermal and material. The intended connection in this paper is therefore a *structural parallel* : in both settings, stability is expressed through minimization of an appropriate functional under constraints.

In the subsections that follow, we will examine the mechanism of U(1) symmetry breaking in full detail. We will also describe its role in the context of superconductivity and the GL free energy density.

### 2.1 Spontaneous Symmetry Breaking of U(1) Gauge Symmetry in Relativistic Ginzburg–Landau (GL) theory

As shown in [27–29], the relativistic GL Lagrangian density can be written in covariant form as^1^

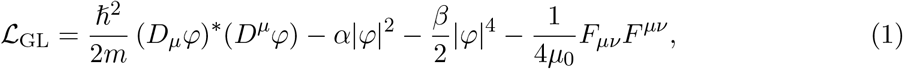

where we use a metric with signature *η*_*µν*_ = diag(−1, +1, +1, +1). Here we identify the following quantities.

- 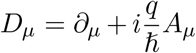 is the gauge covariant derivative, incorporating interactions between the electromagnetic four-potential *A*_*µ*_ and the complex scalar field *φ* consisting of particles with mass *m* and charge *q*.
- *α* and *β* are phenomenological parameters. (In the case of a superconductor, these are determined by the particular material.)
- *F*_*µν*_ = ∂_*µ*_*A*_*ν*_ − ∂_*ν*_*A*_*µ*_ is the electromagnetic field strength tensor.

Gauge transformations of the four-potential and scalar field together with the covariant derivative are given by, respectively,

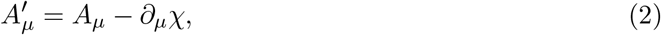

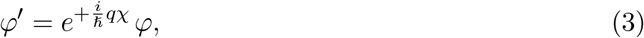

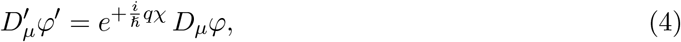

where *χ* is an arbitrary real gauge function. From (1), it is evident that the kinetic term, (*D*^*µ*^*φ*)^∗^ (*D*_*µ*_*φ*), and the full Lagrangian density remain invariant under these gauge transformations. However, the transformation introduces a phase factor *e*^+*iϕ*^ to the scalar field, with 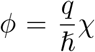. This symmetry corresponds to the global phase rotation of the complex scalar field (for constant *χ*) and describes the global U(1) symmetry.

Noether’s theorem predicts that a global symmetry leads to a conserved current. Using an infinitesimal variation of the field, 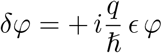 (with *δA*^*μ*^ = 0), the Noether current is

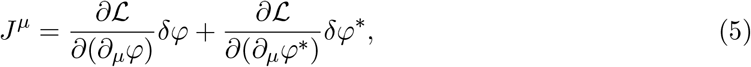

From the Lagrangian density, the four-current density is found to be

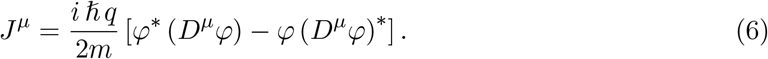

The Euler–Lagrange equation of motion for *A*_*µ*_ is

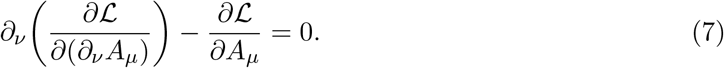

In Lorenz gauge (∂_*µ*_*A*^*µ*^ = 0), this yields

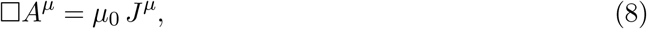

where 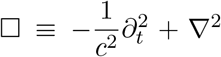. Now we consider the role of spontaneous symmetry breaking of U(1) gauge symmetry. From the Lagrangian density in (1), the potential energy contribution, 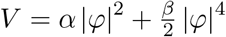, is minimized when

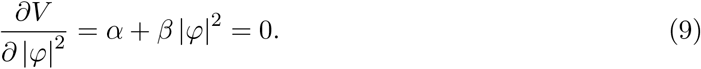

The expression in (9) admits two solutions which are |*φ*|^2^ = 0 (for no symmetry breaking) and 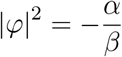 (for the symmetry-breaking phase). Note that 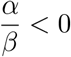 so that |*φ*|^2^ is positive definite in the symmetry-breaking phase.

For the case of no symmetry breaking (|*φ*|^2^ = 0), it is evident from (6) that *J*^*µ*^ = 0. Then (8) becomes simply □*A*^*µ*^ = 0 which leads to the usual wave solution, *A*^*µ*^ ~ *e*^−*i*(*kz*−*ωt*)^, for a massless photon propagating in the *z* direction. However, in the symmetry-breaking phase, 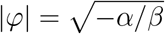 is a constant so 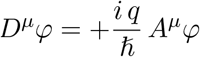. Then (6) becomes

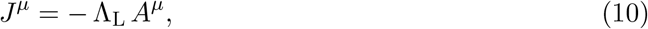

which is the covariant London constitutive equation, where

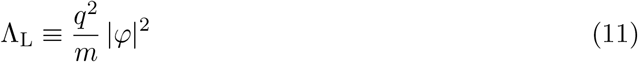

is the London parameter. Consequently, (8) becomes

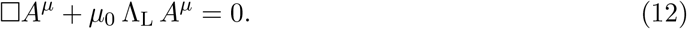

Therefore (12) predicts a solution of the form *A*^*µ*^ ~ *e*^−*z/λ*^. This exponential decay solution implies an attenuation length scale and corresponding mass scale given by, respectively,

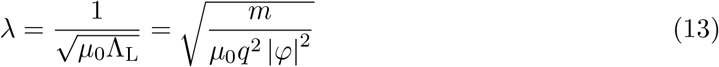

and

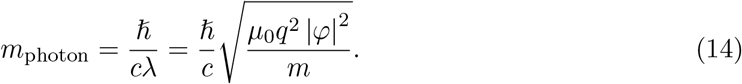

These results demonstrate the Meissner effect for the electromagnetic (EM) field since they show that the EM field is attenuated, or in the zero frequency limit expelled, beyond a length scale given by (13). Also, (14) demonstrates that the photon acquires mass, a well-known feature of U(1) symmetry breaking.^2^

Furthermore, using the symmetry-breaking phase, the Lagrangian density in (1) becomes

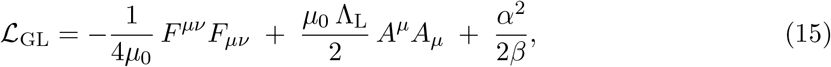

This can be compared to the Proca Lagrangian [28, 29] which describes a massive photon:

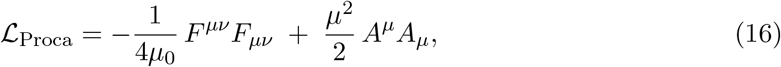

where the photon mass is 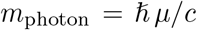. Comparing (15) to (16), it is evident that both contain a mass term involving *A*^*µ*^*A*_*µ*_, and that *µ*^2^ = *µ*_0_ Λ_*L*_. Also note that the last term in (15) is the non-zero ground state energy in the symmetry-breaking phase since it remains even when *A*^*µ*^ = 0. Since the Proca Lagrangian density inherently breaks U(1) gauge symmetry, and the GL Lagrangian density in (1) reduces to the form in (15) which matches the Proca Lagrangian in (16), then the role of U(1) symmetry breaking is confirmed. Furthermore, we have shown the important role of U(1) symmetry breaking as the fundamental mechanism for the Meissner effect. In the next section, we show that superconductivity can be understood as a temperature-dependent spontaneous symmetry breaking which leads to the Meissner effect and London penetration depth associated with superconductivity.

### 2.2 Spontaneous Symmetry Breaking for Superconductivity

The GL model of superconductivity involves a free energy density that describes the superconducting state. We can obtain an energy density from the GL lagrangian density in (1). We begin by evaluating the canonical energy-momentum-stress tensor using

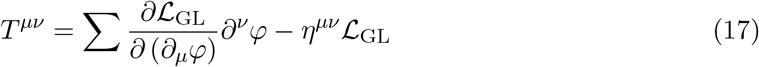

where the sum is over all fields: *φ, φ*^∗^, and *A*_*µ*_. This expression is analogous to the Noether current in (5). In fact, the conserved stress tensor in (17) arises from Noether’s theorem due to space-time translation symmetry. Using (1) and symmetrizing (with respect to indices) leads to^3^

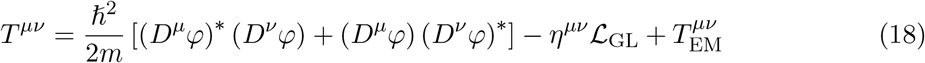

where

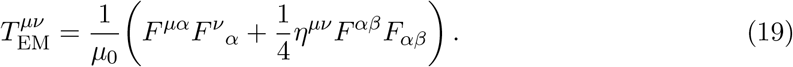

Evaluating *T* ^00^ gives

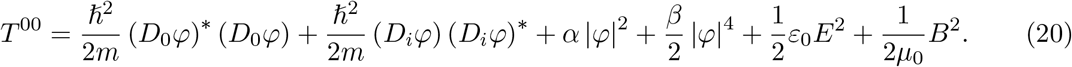

The electric field, 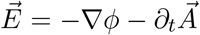, vanishes in the static limit 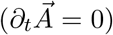 and by using the temporal gauge (*ϕ* = 0), also known as the Weyl or Hamiltonian gauge. Furthermore, using the static limit of the scalar field (∂_*t*_*φ* = 0) leads to

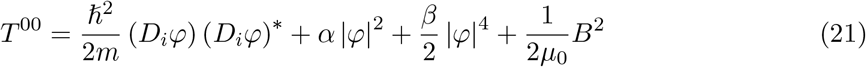

We now have a form that is formally identical to the standard GL free energy density. However, the expression involves a complex scalar field that was initially defined in terms of a relativistic quantum field theory in (1). To justify a reinterpretation of this field in terms of the non-relativistic wavefunction of standard GL theory, recall that a relativistic scalar field (such as described by the Klein-Gordon equation) is related to the non-relativistic Schrödinger wave function by 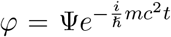. Hence the rest energy simply introduces a constant phase that is invariant under gauge and coordinate transformations.

In the context of the GL model of superconductivity, Ψ is further reinterpreted as a *macroscopic* complex order parameter describing Cooper pairs as a condensate, where |Ψ|^2^ = *n*_s_ is the number density of Cooper pairs. This interpretation as a macroscopic many-particle wavefunction is justified by the Bardeen-Cooper-Schrieffer (BCS) theory [12]. Furthermore, as described in Section 6.3.1 of [31], a gauge transformation of the condensate wave function, 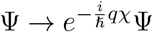, indicates that the macroscopic variable Ψ acquires a phase under a gauge transformation, thereby manifesting symmetry breaking. In this context, the transition from Ψ = 0 (symmetric phase) to Ψ ≠ 0 (broken-symmetry phase) is characteristic of the onset of superconductivity. Furthermore, the GL free energy density can be written using (21) but now in the form

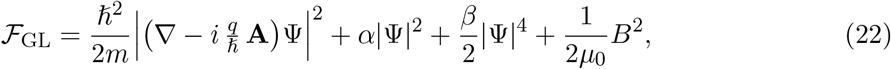

where *q* = −2*e* and *m* = 2*m*_e_ for Cooper pairs. The total free energy of the system is often denoted ℱ = ℱ_*n*_ + ℱ_GL_, where ℱ_*n*_ is the normal state [32]. The GL free energy is minimized by applying the same condition shown in (9). As before, this leads to two solutions given by |Ψ|^2^ = 0 (for no symmetry breaking) and 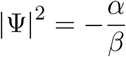 (for the symmetry breaking phase). In the superconducting state, Ψ acquires a non-zero expectation value ⟨Ψ⟩ ≠ 0. Hence the presence of the Cooper pair condensate signifies the spontaneous breaking of the U(1) gauge symmetry, a fundamental mechanism underlying superconductivity. Near the critical temperature *T*_*c*_, the parameters *α* and *β* are approximated as:

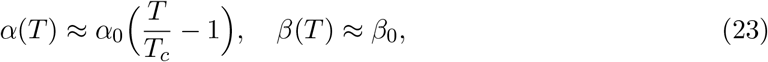

where *α*_0_ and *β*_0_ are positive constants [33]. When *T* < *T*_*c*_, then *α* < 0 which means 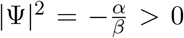, corresponding to a positive Cooper pair density. Conversely, for *T* > *T*_*c*_, we have *α* > 0 which necessitates |Ψ|^2^ = 0, signaling the destruction of the superconducting state. Thus, as the system temperature drops below *T*_*c*_, |Ψ| undergoes a *continuous* (second-order, mean-field) transition from zero to a finite value, embodying the emergence of superconductivity.

Varying the free energy with respect to the vector potential, 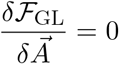, leads to

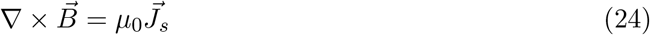

where

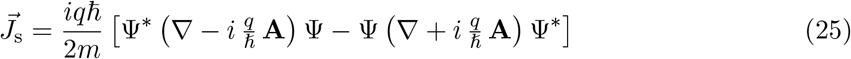

Note that this expression is just the spatial part of the four-current density in (6), as expected. For the case of no symmetry breaking (|Ψ|^2^ = 0), it is evident from (25) that *J*^*i*^ = 0. However, in the symmetry-breaking phase, 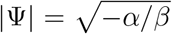 is a constant so ∇Ψ = 0. Then (25) becomes essentially just the spatial part of (10), namely,

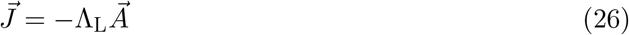

where Λ_L_ is the London parameter given by (11). Note that because |Ψ|^2^ = *n*_s_ is constant, then the charge density of Cooper pairs, *ρ* = *qn*_s_, must also be a constant. Using the continuity equation, 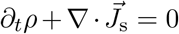 with ∂_*t*_*ρ* = 0, and (26) leads directly to 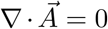 which is the London (Coulomb) gauge. This means the gauge is fixed within a superconductor in equilibrium which is further evidence of gauge symmetry breaking.^4^

Next, we can take the curl of (24) and make use of 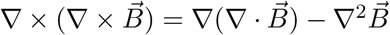 where 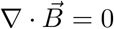. Also using (26) leads to

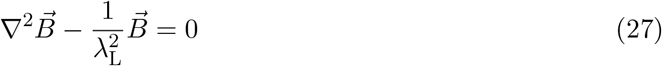

This Yukawa-like equation predicts the well-known Meissner effect for a static magnetic field applied to a superconductor, where *λ*_L_ is the London penetration depth given by (13). The symmetry breaking process underlying these phenomena is illustrated schematically in Fig. 1.

**Figure 1:**
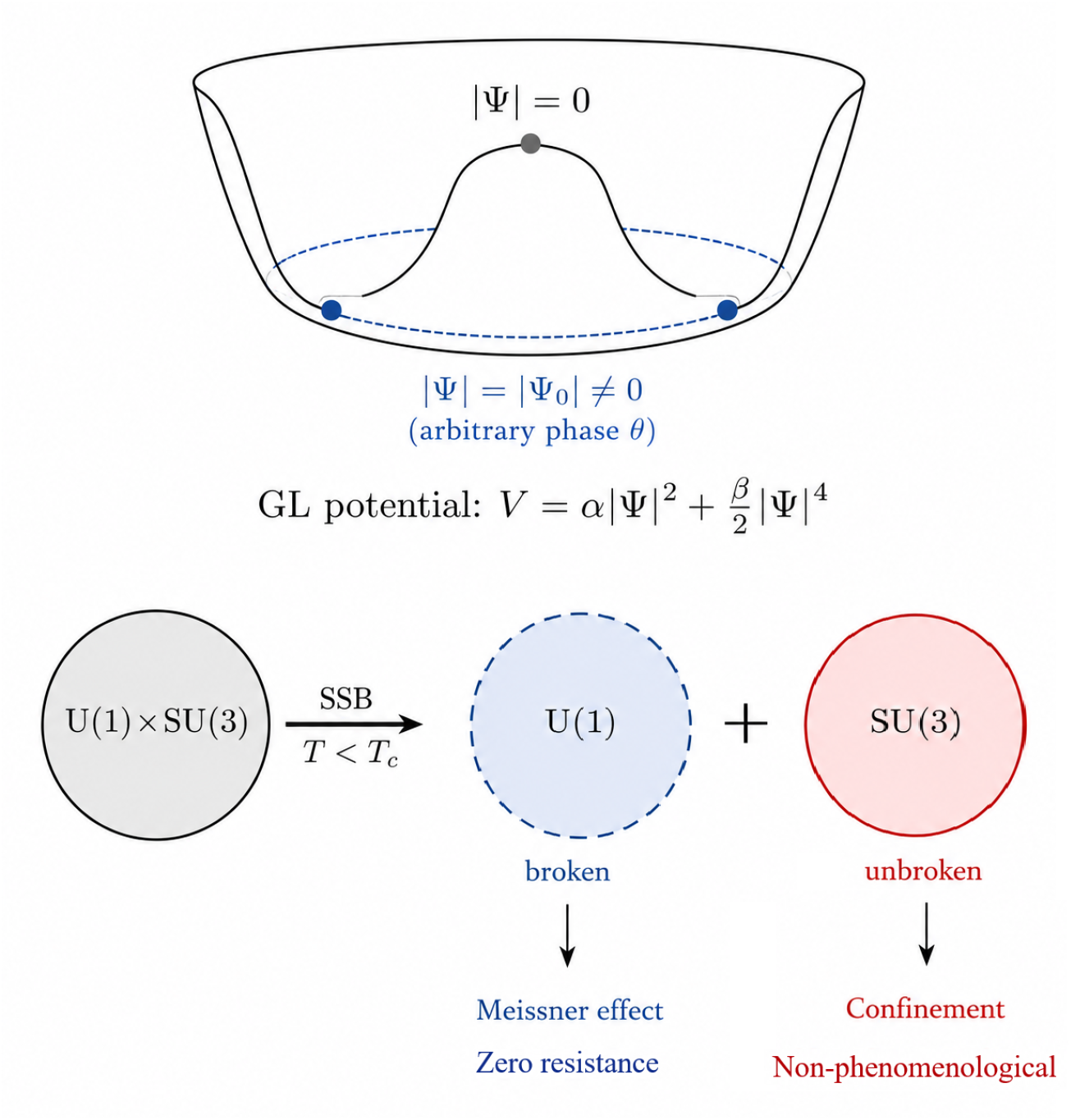
Spontaneous symmetry breaking (SSB) in the superconducting framework. **Top:** The Mexican hat (Ginzburg-Landau) potential shows how the symmetric state (|Ψ| = 0) becomes unstable below *T*_*ϕ*_, and the system falls into a broken-symmetry ground state (|Ψ| ≠ 0). **Bottom:** The symmetry breaking cascade: U(1) electromagnetic symmetry breaks, giving rise to the Meissner effect and zero resistance, while SU(3) remains unbroken, providing the stable foundation for the cosmic vacuum.

### 2.3 Quasi One-Dimensional Superconductivity

In spite of high potential for reducing dissipation and providing global coherence, not all types of superconductivity suit well to biological systems. In particular, three-dimensional (3D) superconductivity, widely investigated in condensed matter physics, is too rigid for living organisms not giving them enough freedom and flexibility for adaptations. 2D, and even in higher degree 1D, superconductivity suits biological systems much better.

The main difference between 3D and lower-dimension types of superconductivity is that in 3D the characteristic temperatures are almost the same: *T*_pair_, where Cooper pairs form and the superconducting gap opens; *T*_*ϕ*_, where global phase coherence appears; and *T*_*R*=0_, where measured resistance vanishes. In reduced dimensions, they can be dramatically different. For example, in 2D, *T*_*ϕ*_ (often a Berezinskii–Kosterlitz–Thouless coherence scale) can be considerably lower than *T*_pair_; in that case true zero resistance on macroscopic scales is expected only at (or below) *T*_*ϕ*_ rather than at *T*_pair_. In 1D, *T*_*ϕ*_ is much lower than *T*_pair_, and *T*_*R*=0_ is typically zero due to phase slips. In quasi-1D superconductors, inter-channel coupling can render *T*_*R*=0_ finite, and it becomes important how far it lies below *T*_*ϕ*_.

Since in biological systems superconductivity comes from microtubules, which are filamentary structures, quasi-1D superconductivity is very natural for living organisms. In this section, we calculate critical temperatures and important parameters for biological superconductors based on available experimental data.

*T*_pair_ was inferred by Mikheenko from an interpreted gap-like feature in [1, 2]. Its reported value is 2022 ± 157 K. This critical temperature is defined by the local pairing energy, and it is large because the pairing is local electronic pairing shielded from thermal decoherence rather than phonon-mediated. It is not an exotic characteristic temperature. Excitons in organic compounds as well as polarons and even hydrogen bond networks have similar or higher characteristic temperatures. Below *T*_pair_, however, coherence in the superconducting condensate is absent down to *T*_*ϕ*_. Between these temperatures, coherence collapses because the long-range phase order is destroyed by fluctuations; thermal phase slips are then dominant in the behavior of the condensate.

Since *T*_*ϕ*_ is the characteristic temperature for the appearance of coherence, it is of primary importance for living organisms. The usefulness of *T*_pair_ and the pairing mechanism can be reduced to providing finite superconducting carrier density *n*_*s*_, which defines the value of *T*_*ϕ*_. In a 1D system, for the derivation of *T*_*ϕ*_, *n*_*s*_ needs to be converted to 1D linear density *λ* = *n*_*s*_ · *A*, where *A* is the cross-section of the superconducting channel, and the concept of superconducting phase stiffness needs to be introduced.

The simplest way to do this is to calculate the energy needed to twist the superconducting phase of the condensate by one radian. This energy depends on the length *L* of the superconducting segment under consideration:

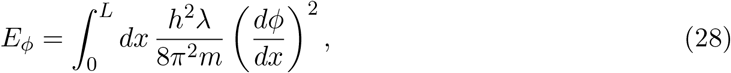

where *m* is the mass of superconducting charge carriers (double the electron mass), and *h* is Planck’s constant. In Eq. (28),

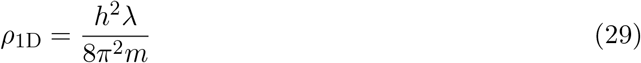

is the 1D stiffness parameter. Its units are [energy · length].

The energy cost of twisting the phase by one radian over a segment of finite length *L* is then *ρ*_1D_*/L*. By equating it to *k*_*B*_*T*_*ϕ*_, where *k*_*B*_ is the Boltzmann constant, the estimate for *T*_*ϕ*_ is:

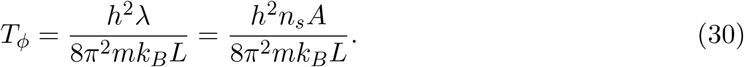

Assuming that superconductivity develops in the lumen of a microtubule with cross-section *A* = 1.77 × 10^−16^ m^2^, for an essentially metallic superconducting density of *n*_*s*_ ≈ 8 × 10^27^ m^−3^, and a segment length of 1 micrometer, *T*_*ϕ*_ = 310 K. These choices are calibrated to the reported measurements but remain model assumptions. In Fig. 2, the screening of magnetic flux by the cross-sections of microtubules imaged by magnetic force microscopy is shown. In a), the topographic image displays the lumens of several microtubules (dark holes), while b) shows a magnetic response, with bright areas corresponding to the expelled magnetic flux. In c), the high-resolution image of expelled flux of an individual microtubule is shown. The flux is expelled from all of the lumen.

**Figure 2:**
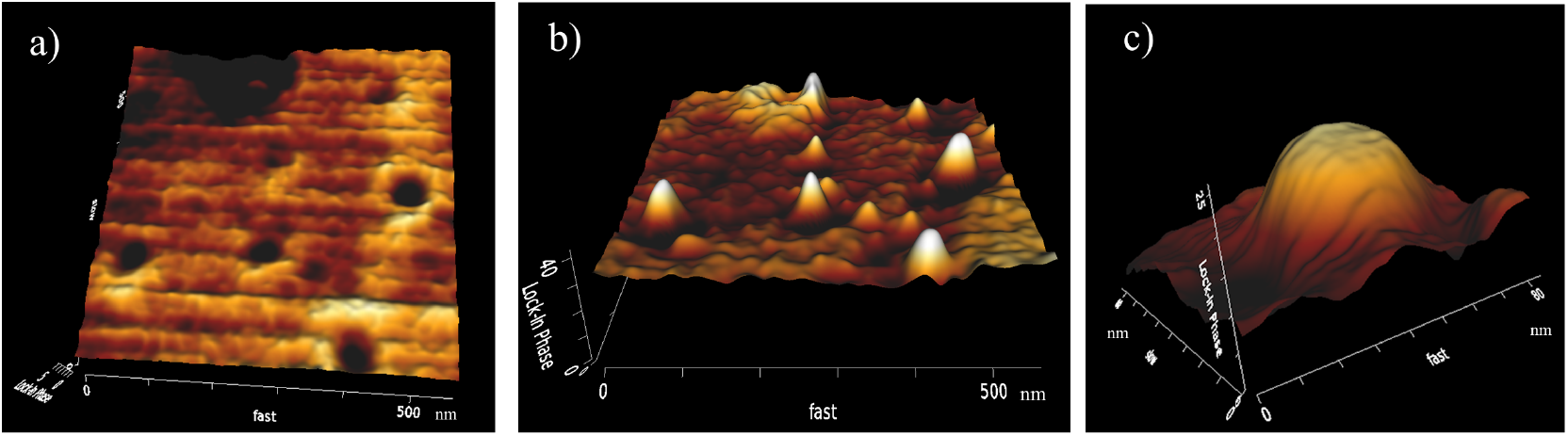
Screening of magnetic flux by the cross-sections of microtubules imaged by magnetic force microscopy. In a), the topographic image displays the lumens of several microtubules (dark holes), while b) shows a magnetic response, with bright areas corresponding to the expelled magnetic flux. In c), the high-resolution image of expelled flux of an individual microtubule is shown. The flux is expelled from all of the lumen. Adapted from the reported magnetic-force-microscopy work in Refs. [1, 37]; permission or licence status should be confirmed before publication.

In Fig. 3, the in-plain growth of the microtubules from a drop of tubulin protein in the top-right corner is shown. Image b) is recorded 10 minutes after image a), and image c) is recorded 6 minutes after image b). The bright color corresponds, as in Fig. 2, to the superconducting state, while the dark color corresponds to the normal state. Most microtubules are superconducting, but some remain normal and are growing as normal. Moreover, some segments of microtubules, compare c) with a) and b) in the area not far from the center, switch fully from the superconducting to normal state. The length of either superconducting or normal segments is close to 1 micrometer.

**Figure 3:**
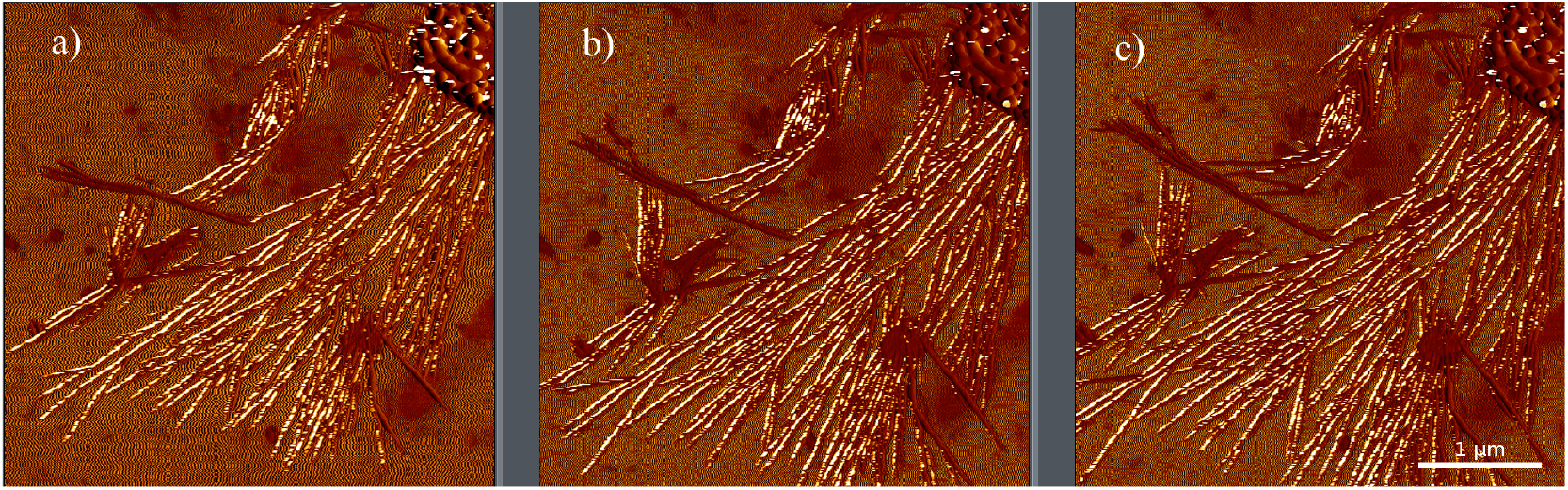
In-plain growth of the microtubules from a drop of tubulin protein in the top-right corner. Image b) is recorded 10 minutes after image a), and image c) is recorded 6 minutes after image b). The bright color corresponds, as in Fig. 2, to the superconducting state, while the dark color corresponds to the normal state. Most microtubules are superconducting, but some remain normal and are growing as normal. Moreover, some segments of microtubules, compare c) with a) and b) in the area not far from the center, switch fully from the superconducting to normal state. The length of either superconducting or normal segments is close to 1 micrometer. Adapted from the reported microtubule-imaging work in Refs. [1, 37]; permission or licence status should be confirmed before publication.

Such a requirement for *n*_*s*_ could be relaxed if superconductivity develops in the whole cross-section of the microtubule plus a thin layer of structured water around it. Alternatively, coherence could spread to longer fragments of the microtubules.

Using the mentioned *n*_*s*_ ≈ 10^28^ m^−3^, the Fermi velocity *v*_*F*_ for the charge carriers can be found, and the coherence length of the superconductor can be calculated using the relation:

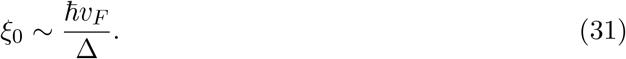

This gives a coherence length for the biological superconductor on the order of 1 nm, as reported in [1]. Moreover, the value of 10^28^ m^−3^ allows calculation of the magnetic penetration depth using:

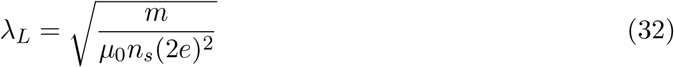

where *µ*_0_ is the magnetic permeability of vacuum and *e* is the electron charge. The calculation gives *λ*_*L*_ ≈ 38 nm, which is also close to the experimental value reported in [1]. This internal numerical consistency motivates further testing of the proposed 1D superconductivity interpretation in microtubules. In spite of the very high *T*_pair_ of 2022 K, *T*_*ϕ*_ is reasonably low. Moreover, it is close to the temperatures at which animals stabilize their bodies. This physiological-scale proximity is suggestive within the model, but it is not independent evidence for evolutionary optimization.

Physically, with establishment of coherence below *T*_*ϕ*_, resistance starts exponentially decreasing to zero, as shown schematically in Fig. 5. Within this model, the physiological relevance of *T*_*ϕ*_ is a hypothesis: operation near this threshold could preserve both coherence and adaptability, whereas operation far below or above it would be expected to reduce flexibility or coherence, respectively.

Being close to 310 K is also important because this temperature lies within the physiological range in which microtubules remain dynamic. The detailed temperature windows for microtubule polymerization, depolymerization, protofilament peeling, and tubulin denaturation are preparation-dependent; they are used here only as biological context for the model, not as independent evidence for the superconducting threshold.

Assuming that microtubules are in the correct temperature range, one can ask how resistance would drop below *T*_*ϕ*_ and whether *T*_*R*=0_ would be achieved. The answer depends on the number of microtubules in a neuron, their packing in bundles and the strength of superconducting correlations between them. The numerical separation between *T*_*ϕ*_ and *T*_*R*=0_ is therefore treated as a model-dependent estimate requiring direct derivation or experiment, rather than as an established biological value. The microtubule structure and the relevant temperature scales are illustrated in Fig. 4.

**Figure 4:**
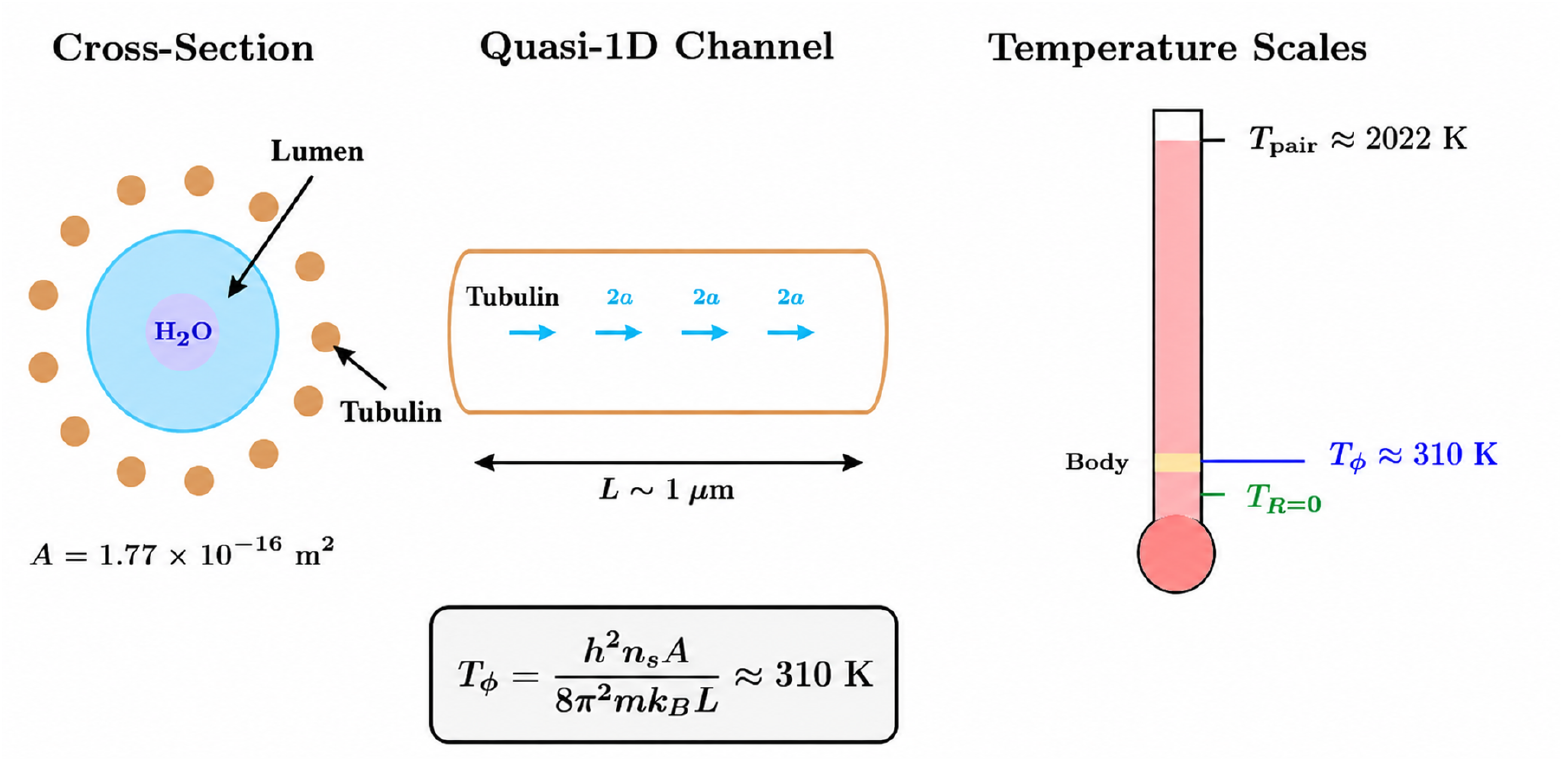
Microtubule structure and quasi-1D superconductivity. **Left:** Cross-section showing the tubulin dimer ring surrounding the water-filled lumen where superconducting charge carriers flow. **Center:** Longitudinal view of the quasi-1D superconducting channel with Cooper pairs (2e). **Right:** Temperature hierarchy showing the pairing temperature *T*_pair_ ≈ 2022 K and the coherence temperature *T*_*ϕ*_ ≈ 310 K, with body temperature schematically positioned at this threshold.

**Figure 5:**
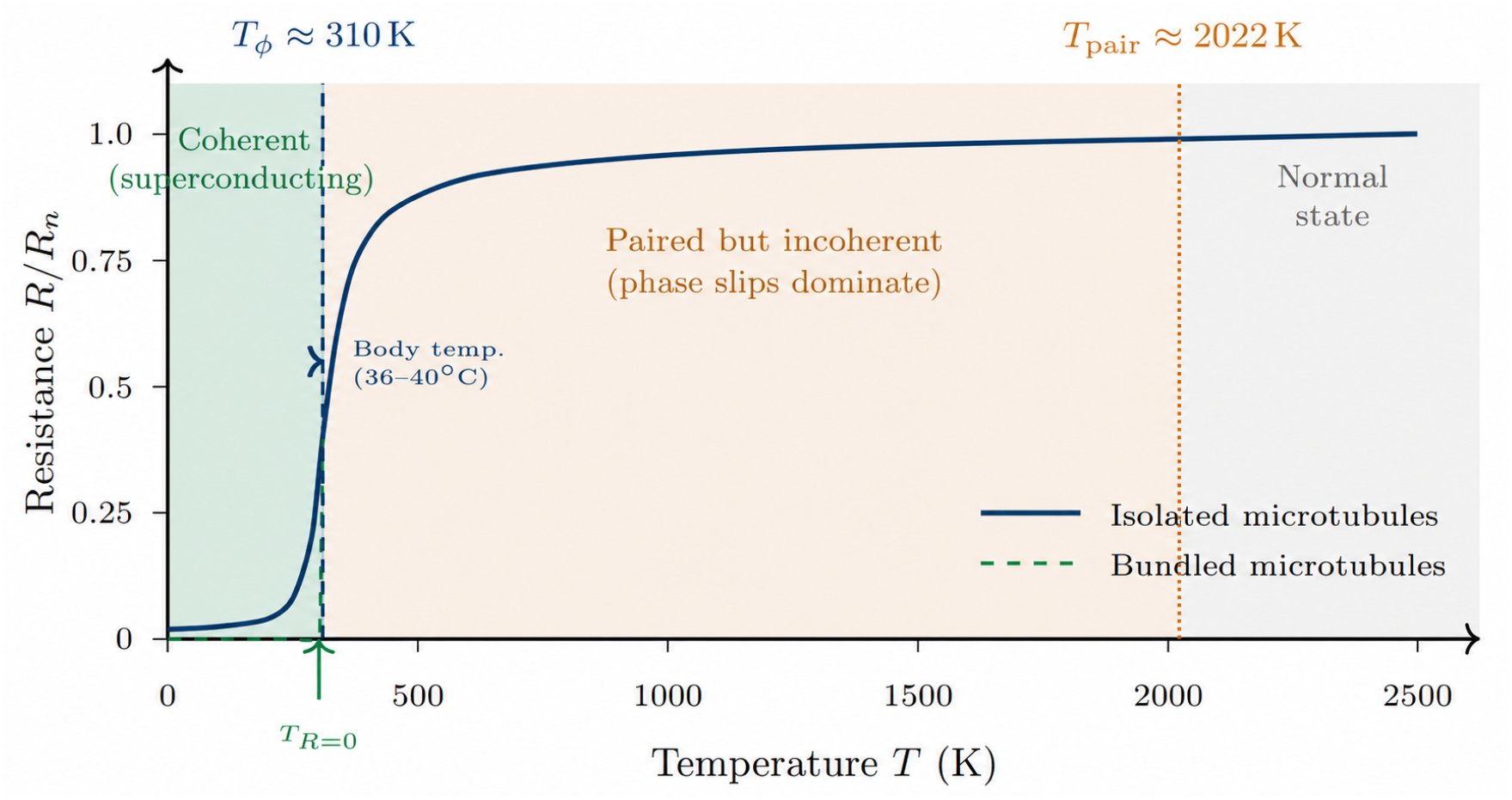
Schematic resistance *R*(*T*) for quasi-1D biological superconductivity. Three regimes are shown: (I) coherent superconducting state below *T*_*ϕ*_ ≈ 310 K where resistance drops exponentially; (II) paired but incoherent regime between *T*_*ϕ*_ and *T*_pair_ ≈ 2022 K where Cooper pairs exist but phase coherence is destroyed by thermal fluctuations; (III) normal state above *T*_pair_. The solid blue curve represents isolated microtubules with residual resistance from phase slips; the dashed green curve shows tightly-bundled microtubules achieving true zero resistance (*T*_*R*=0_) within ~1 K of *T*_*ϕ*_. The shaded band marks physiological body temperature (36–40^◦^C), positioned near the coherence threshold within the schematic model.

## 3 Implications for Brain Superconductivity and Consciousness

If biological systems such as the brain can support superconducting-like behavior near physiological conditions, this would open a new way to examine consciousness through quantum coherence. Motivated by recent experimental reports of superconductivity-like phenomena in neuronal microtubules, we extend the Ginzburg–Landau (GL) description of superconductivity to neural dynamics. We hypothesize that some neural states may access superconducting-like regimes in which effective dissipation is strongly suppressed relative to ordinary matter, allowing enhanced coherence to persist. If this assumption is confirmed, it could offer a physical route to two long-standing problems in brain science and consciousness: binding and neural efficiency.

Stated conservatively, superconductivity offers a candidate mechanism for organizing non-separable large-scale structures under noisy conditions, and binding requires widely distributed neural populations to behave, at least transiently, as a single coordinated state. This requirement is not naturally met by message passing alone between separate component systems. It is more naturally expressed through a common order parameter that can include many degrees of freedom within one coordinated phase. Coherence is therefore treated here as a possible binding resource, recruited when binding is both needed and physically available.

The energetic cost of long-range integration need not be accounted for only by noisy dissipation. In coherent phases of organization, part of this cost may be carried by phase organization across a large collective structure. This allows a phase- and state-dependent equilibrium in which organization increases when collective processing is needed, and relaxes again when attention shifts toward local plasticity or flexible processing. Therefore, ordinary cognition and altered states of experience may differ by the relative balance between active dissipative processing and state-dependent pairing.

It is interesting to consider how the two-fluid aspects of the brain under the present model map onto conscious and non-conscious states. As shown in Fig. 6, we connect the approach to a superconducting state with Metzinger’s concept of minimal phenomenal experience (MPE) [38]. We treat *content-minimal absorption* as an experiential state in which reportable content becomes weak, while attention and alertness remain stable. Such states may reflect a shift in large-scale neural coordination towards a coherence threshold. We treat meditative cessation—the complete absence of consciousness—differently. Cessation marks a non-phenomenological interval beyond ordinary experience, where reportable contents disappear but a latent organizing structure persists [39, 40].

**Figure 6:**
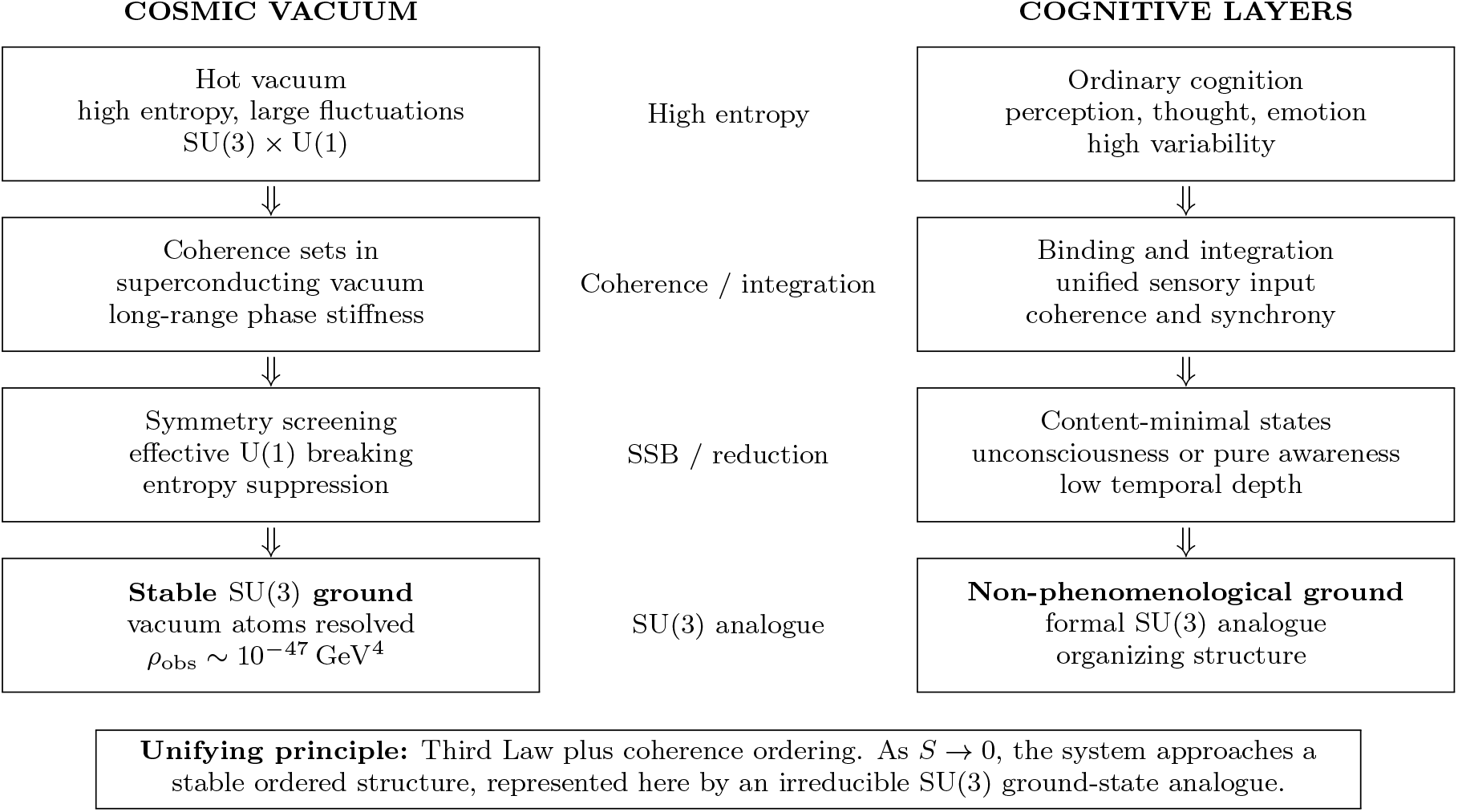
A layered analogy between the cosmic-vacuum construction and the cognitive model. In the SU(3) vacuum-atom framework, *ρ*_vac_ is tied to the realized low-entropy vacuum state. In cognition, the right column is only a structural analogy: integration supports unified contents, reduced ordinary organization corresponds to content-minimal or low-complexity states, and the final SU(3) node is a non-phenomenological organizing analogue rather than particular experienced content.

In the superconductivity analogy we adopt:

- The superconducting parameter Ψ, from the Ginzburg–Landau theory, corresponds to the emergent coherent neural field formed by synchronized neuronal oscillations.
- The spontaneous symmetry breaking of the electromagnetic U(1) gauge symmetry reflects the neural transition into a more unified state. In the cross-domain framework developed here, this is paired with a *separate* structural hypothesis: namely, that a stable non-Abelian organizing residue may be modeled by SU(3). In the neuroscience application, SU(3) functions as a latent non-factorizing geometry that constrains how reportable conscious composites can emerge and bind. The observable phenomenological level corresponds to such composites, while the underlying SU(3) degrees of freedom enter through inferred invariant structure.

This approach also addresses a common objection to quantum theories of consciousness, including the Orch-OR model of Hameroff and Penrose [41]. The main concern is decoherence: at biological temperatures, ordinary quantum states are expected to lose coherence rapidly, as emphasized in direct decoherence critiques of brain quantum models [42]. Our proposal takes a different route. It asks whether structured water and confined channels inside neuronal microtubules can support superconducting-like states that protect coherence more effectively than ordinary biological matter. In this setting, the relevant coherence time, *τ*_*c*_, may be estimated as:

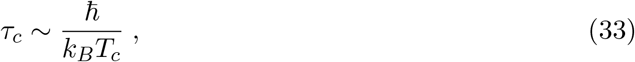

Here, 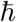 is the reduced Planck constant, *k*_*B*_ is Boltzmann’s constant, and *T*_*c*_ is the critical temperature of the proposed superconducting-like state. The estimate gives only a crude microscopic thermal timescale. It should not be treated as a direct measurement of neural phase coherence. If biological superconductivity is experimentally confirmed and a measurable *T*_*c*_ is established, the associated coherence times can then be tested against neurophysiological data.

The two-fluid superconductivity model, originally developed for describing superconductors at a microscopic level, provides an insightful parallel for neural dynamics. Under this model, neural activity may simultaneously involve a classical (dissipative) neural component coexisting with a quantum-coherent (dissipationless) superconductive component. Phenomenologically, this coexistence predicts that some high-integration states could show improved energetic efficiency relative to comparable purely dissipative dynamics; this should be tested with metabolic or neurovascular measurements rather than assumed.

The brain organizes many local neural states into coherent wholes. This fits naturally with hierarchical self-organization in the Free Energy Principle. Wiese [43] argues that neural computation may be necessary for consciousness, but it may not be sufficient unless the system also satisfies specific implementation conditions. Ramstead *et al*. [44] develop a related idea through the inner screen hypothesis, where nested Markov blankets help organize neural dynamics into globally coherent states. In our superconductivity analogy, neural symmetry breaking supplies a possible physical implementation condition. It provides a mechanism by which low-entropy coherent neural states can become stabilized.

Recent experimental work supports the motivation for this hypothesis. Belli *et al*. [14] report a strong correlation between bonding-network descriptors and predicted critical temperature in hydrogen-based superconductors. Mikheenko [37] reports superconductivity-like flux expulsion in self-assembled microtubules under near-physiological conditions. These reported, not yet independently replicated, results motivate the possibility that living systems may host superconducting or superconducting-like organization, and that such organization may provide a physical basis for quantum coherence in neural matter. This coherence fits naturally with the Free Energy Principle, since both emphasize stable organization under constraints. In this framework, life may exploit superconductive-like order to preserve coherent structure despite biological noise, while also providing a frictionless, energy-efficient way to organize experience into coherent wholes. Conscious states can then be understood as emergent states of a highly coherent neural system. This view connects with the criterion for conscious organisms discussed by Laukkonen *et al*. [45]: a stable, unified reality model with internal organization capable of supporting sustained integration and reflexivity.

As discussed earlier, Mikheenko [2] reported unexpected phenomena in brain tissue, including an abrupt drop in resistance, evidence for an energy gap, and Josephson-like electro-magnetic emission. These signals indicate an extremely high inferred critical temperature of approximately 2022±157 K, but they require independent verification. If confirmed, they would strengthen the analogy with Little’s quasi-one-dimensional organic superconductors and suggest a possible physical basis for rapid neural information transfer. The idea separates local pairing from global coherence. Local pairing may occur in microtubules or structured water without implying global superconductivity throughout the brain. A phase-ordering process could then allow many local elements to act as a cooperative whole. In this picture, coherent phase ordering acts as a binding amplifier, increasing or decreasing its effect according to the processing demands of the system.

The brain already contains ordinary biological controls that could move it closer to, or further from, a coherence threshold. These include local temperature variations, metabolic rate, hydration, tonicity, microtubule polymerization, ionic composition of the extracellular fluid, and neuromodulators that influence cytoskeletal and membrane dynamics. The brain already uses such parameters to adjust excitability and synchrony. A coherent channel, if present, would likely be coupled to these same biological adjustments. Any practice that changes global neural synchrony may therefore also shift the system toward or away from this threshold.

Empirically, altered-state dynamics need not manifest as a monotonic increase in classical coherence measures. In particular, advanced absorption has been reported to shift brain dynamics toward a metastable, near-critical regime with increased signal diversity and reorganized long-range temporal correlations [46]. This motivates a more conservative prediction style: the model predicts state-dependent reorganization across impedance/coupling structure and near-critical dynamics as the system is tuned toward or away from a coherence threshold. In subsequent sections, we develop and mathematically formalize these conceptual connections within the SU(3)-based neural geometrodynamics framework.

## 4 Two-fluid Model: Evidence from Quantum and Electromagnetic Field Interactions in the Brain

The two-fluid model is a cornerstone of superconductivity theory. It describes how a superconducting material supports two distinct types of charge carriers simultaneously:

- *Superfluid Component* 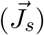: Represents the Cooper pair condensate that flows without electrical resistance, carrying the dissipationless supercurrent.
- *Normal Fluid Component* 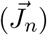: Comprises unpaired electrons or quasiparticles that obey Ohm’s law and contribute to non-zero electrical resistance.

The application of this model to neural states is captured by how the balance between dissipative and coherent components can shift across cognitive regimes and physiological conditions.Mathematically, the total current density is given by:

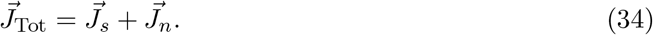

In conventional superconductors, one typically writes:

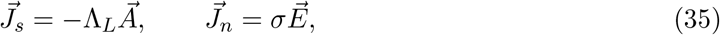

where *σ* is the Ohmic conductivity.

### 4.1 Application to Neural Dynamics

It is important to clarify that “coherence” in neural dynamics is often quantified by phase synchronization between signals recorded from different brain regions. In classical superconductors, however, coherence implies a macroscopic phase-locking that results in dissipationless transport. In our model, altered states are treated as possible regimes in which neural coherence may play a role analogous to macroscopic phase coherence, without assuming that ordinary neural synchrony is itself superconductivity. In practical terms, this would mean that, when the brain enters a coherent state 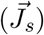, measurable quantities such as brain criticality, together with diminished effective resistance, become pronounced, signaling a transition toward superconductive-like behavior. The resulting claim is a prediction of the two-fluid model: if such a phase exists, it should reduce effective dissipation and improve information transfer relative to dissipative alternatives.

We define a neural order parameter *ψ*(**r**, *t*) whose squared magnitude represents the local density of neurons engaged in coherent (supercurrent-like) activity:

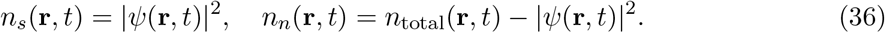

Thus, the total neural current can be written as

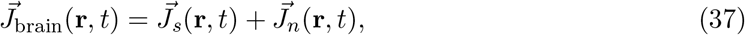

with 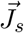 proportional to |*ψ*(**r**, *t*)|^2^ and 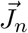 representing the dissipative component.

### 4.2 Dynamic Transition Between Neural States

The transition from a fully conscious to a meditative state may be conceptualized hypothetically as a phase transition analogous to that observed in superconductors:

- *Initiation of Meditation*: In the model mapping, partial symmetry breaking in neural circuits corresponds to Cooper pair–like assembly formation and increased coherence.
- *Intermediate State*: The brain exhibits a mixed state, with both coherent (superfluid-like) and incoherent (normal) neural activity coexisting.
- *Deep Meditative State*: Within the same hypothetical mapping, if the effective neural temperature *T*_eff_(*t*) falls below a critical threshold *T*_*ϕ*_, the order parameter *ψ*(**r**, *t*) grows, leading to a dominant superfluid-like state with minimal resistance.

We introduce an effective neural temperature *T*_eff_(*t*) such that:

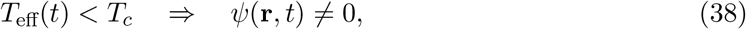

and conversely, if *T*_eff_(*t*) > *T*_*ϕ*_, then *ψ*(**r**, *t*) = 0. This is a modeling analogy drawn from the time-dependent Ginzburg–Landau (TDGL) formalism, not an empirical claim that meditation has already been shown to cross a superconducting threshold. In that context, the TDGL equation is given by

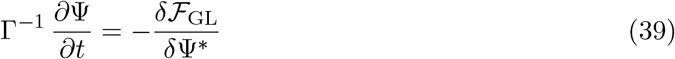

where the relaxation rate Γ governs the dynamics of Ψ.

### 4.3 Implications for Electrical Resistance and Cognitive Function

The two-fluid model applied to neural dynamics yields several significant predictions regarding both the electrical properties of brain tissue and its impact on cognitive function. In our framework, the brain is conceptualized as comprising two interacting components: a coherent (supercurrent-like state) and a dissipative (normal current-like) state. This dualistic model leads to the following key implications:

- *Reduction in Electrical Resistance:* As the proportion of the coherent, superconducting-like component increases, the overall electrical resistance in neural circuits is expected to decrease. Such a reduction facilitates faster, more efficient signal propagation, analogous to the zero-resistance state observed in conventional superconductors.
- *Enhanced Energy Efficiency:* Increased neural coherence minimizes energy losses due to dissipation. This reduction in energy expenditure may lower metabolic demands and allow the brain to maintain prolonged periods of efficient information processing.
- *Improved Information Integration:* High levels of neural coherence foster stable global synchronization across brain regions, which is critical for integrated cognitive processing. Such unified neural activity may underlie the minimal phenomenal experiences (MPE) described by Metzinger [38], and is likely to be observed during deep meditative or psychedelic states.

One can express a schematic frequency-dependent neural resistivity interpolation as

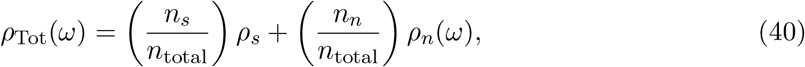

where *ρ*_*s*_ = 0 denotes the near-zero resistivity of the superconducting-like (coherent) component, *n*_*s*_ is its density, *n*_*n*_ is the density of the normal (dissipative) component, and *n*_total_ = *n*_*s*_ + *n*_*n*_. As the ratio *n*_*s*_*/n*_total_ increases, the overall resistivity *ρ*_Tot_(*ω*) approaches zero, mirroring the behavior observed in conventional superconductors. This equation is used only as a phenomenological interpolation; for parallel transport channels, conductivities or admittances should be combined instead.

The Cole–Cole law was introduced as an empirical dielectric-dispersion model, while brain-tissue electrical parameters require tissue-specific conductivity sources [47–49]. As an illustrative normal-component model, we write

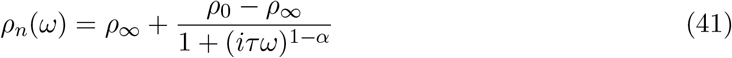

where *ρ*_0_ is the resistance at 0 Hz (DC), *ρ*_∞_ is the resistance for *ω* → ∞, *τ* is the time constant related to membrane charging, and *α* is the dispersion parameter. For illustration only, one may take *ρ*_0_ ~ 10^3^ Ω·cm, *ρ*_∞_ ~ 2 × 10^2^ Ω·cm, *τ* ~ 10^−4^ s, and *α* ≈ 0.7. Therefore (41) becomes

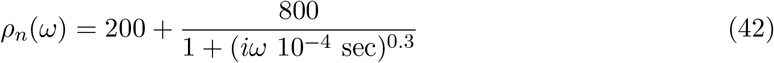

in units of Ω·cm. A plot of this function appears in Fig. 7 below. Note that substituting (41) into (40), and using *ρ*_*s*_ = 0 and *n*_total_ = *n*_*s*_ + *n*_*n*_, yields

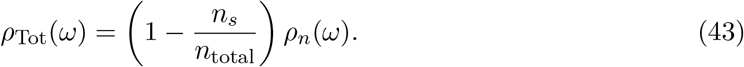

**Figure 7:**
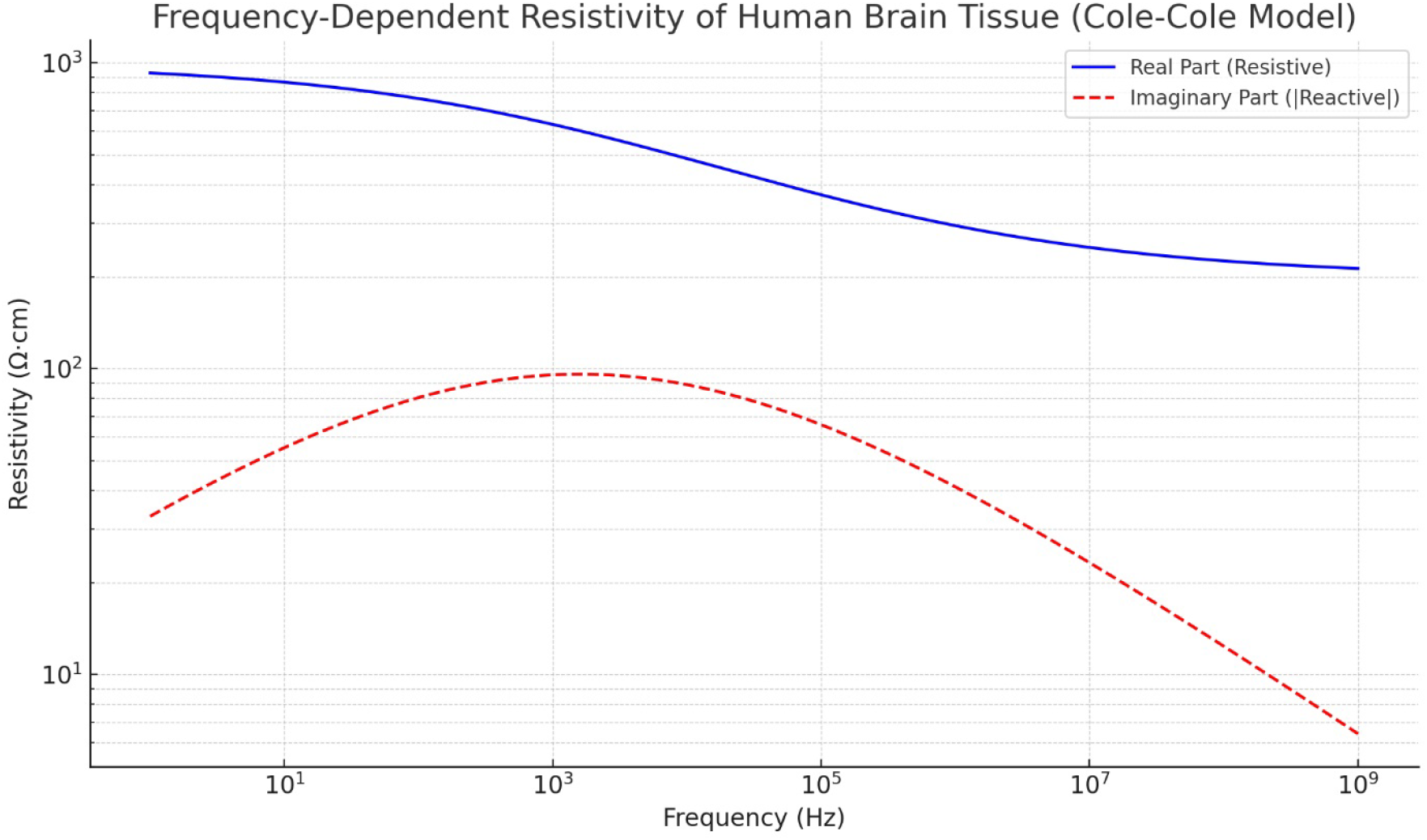
Illustrative plot of resistivity (*ρ*_*n*_) versus frequency (*ω*) for a Cole–Cole-type normal-component model applied to brain tissue.

This expression demonstrates that for a given frequency, the plot of *ρ*_Tot_ is expected to shift down relative to *ρ*_*n*_ predicted by the Cole-Cole model. The amount of this shift is given by *n*_*s*_*/n*_total_ which becomes larger as the brain enters into a deeper superconductive state.

Furthermore, unlike in conventional superconductors, where only cooling below the material’s critical temperature *T*_*c*_ induces the coherent phase, psychophysiological studies motivate the possibility that contemplative disciplines can steer the *effective* neural temperature *T*_eff_. Benson *et al*. reported large peripheral warming during Tibetan *g Tum-mo* without a rectal-temperature increase [50], while Kozhevnikov *et al*. reported axillary-temperature increases during forceful practice, with visualization helping sustain them [51]. The upper-respiratory cooling study of Mariak *et al*. concerned neurosurgical patients with an upper-airway bypass, not slow nasal or diaphragmatic breathing [52]. In our framework, such thermal observations are treated only as context for *T*_eff_; once this variable crosses the fixed material threshold *T*_*ϕ*_ (set by the GL coefficients *α, β*), a non-zero condensate fraction *n*_*s*_ can, in principle, emerge without any need to modify the intrinsic superconducting constants themselves.

Moreover, studies such as those by Tidswell et al. [53] using three-dimensional Electrical Impedance Tomography (EIT) demonstrated feasibility for detecting activity-related impedance changes from scalp measurements, although early reconstructions were technically limited. The further claim that a superconducting-like transition could enhance signal propagation or cognitive efficiency is a prediction of the present model, not a conclusion established by EIT alone.

### 4.4 Experimental Predictions and Testing

To assess the relevance of the two-fluid picture in neural systems, the experimental program should remain cautious and indirect. The more defensible strategy is to search for convergent proxies of state-dependent coordination and coherence-threshold efficiency:

- *Large-scale coordination proxies*: Use EEG, MEG, or intracranial recordings to examine changes in phase locking, long-range temporal correlations, metastability, or criticality-like signatures across altered states.
- *State-dependent coherence scales*: Estimate empirical coherence lengths and times from neural oscillations and autocorrelation structure, and compare them with the model’s predicted shifts near a coherence threshold.
- *Energetic efficiency*: Combine electrophysiology with metabolic or hemodynamic measurements to test whether certain high-integration states are accompanied by reduced effective energetic cost.
- *Thermal and microstructural context*: Use MR thermography or related methods, where feasible, together with measures sensitive to cytoskeletal or hydration state, to investigate whether ordinary physiological variables modulate access to coherence-supporting regimes.

TDGL-inspired neural modeling may still be useful as a phenomenological scaffold, but its predictions should be framed gently. The most realistic near-term goal is to test whether the coupled dissipative/coherence-supporting description organizes data better than purely dissipative alternatives. In that spirit, the model predicts state-dependent shifts in integrative coordination across altered states.

By adopting a two-fluid model, we conceptualize brain dynamics as arising from a partitioning of neural activity into a coherence-supporting component and a dissipative component. Using a time-dependent order parameter *ψ*(**r**, *t*) and the corresponding TDGL formalism, we can model transitions in the relative dominance of these components as the effective neural temperature *T*_eff_(*t*) moves relative to a critical threshold *T*_*ϕ*_. This framework motivates testable signatures in coordinated variability and energetic efficiency while remaining compatible with the possibility that some deep or content-minimal states involve reduced ordinary large-scale coherence rather than maximal synchrony.

## 5 It from SU(3)

### 5.1 Cosmic Superconductivity: The Vacuum as a Protected Medium

Two hard problems raise a similar structural question. The first comes from cosmology. Quantum field theory assigns an immense natural energy density to the vacuum, while the measured dark-energy scale is smaller by about 10^123^. The second comes from experience. Certain cessation states, reported in advanced meditation practice and recognized in some contemplative traditions, appear to involve the loss of ordinary experience, including the usual sense of self. Reports describe a sudden discontinuity. When descriptions are available, they usually concern the states before and after the interval, rather than any content within the interval itself [39, 40]. Although these problems belong to different fields, they raise the same basic question: what prevents a ground state from becoming unstable, and what structure remains when ordinary fluctuations are driven close to zero?

When a system reaches a low-entropy regime, it does not end in emptiness. It settles into a residual irreducible structure. Ref. [54] treats the vacuum as a superconducting-like medium, with cosmic acceleration acting as a macroscopic analogue of Meissner expulsion. Coherence then reorganizes the vacuum ground state and suppresses certain long-wavelength modes. Building on this picture, Refs. [55, 56] propose that, in the extreme low-temperature vacuum, the electromagnetic sector loses its role as the dominant long-range organizing order. Electromagnetism remains an observed interaction, but it no longer defines the vacuum order parameter. The remaining structure is an unbroken SU(3) sector, selected by confinement physics as a stable non-Abelian residue at low energies.

This structure with an unbroken SU(3) symmetry is protected by third-law of thermodyanmics and can realized through confinement-scale vacuum units [55, 56] that are proposed to account for the cosmological-constant value and the values of nature’s constants. In the brain, we suggest that the same organizing idea may appear in reconstructed collective dynamics. As electromagnetic fluctuation-driven activity is reduced, the remaining integrated organization may take the form of a localized, non-factorizing SU(3) geometry. When fluctuation-generating degrees of freedom are removed, suppressed, or made dynamically unavailable, the stable ground regime is organized by an irreducible SU(3) structure. A strength of this approach is that both sides appeal to irreducibility. Theories of consciousness treat experience as unified organization. Integrated Information Theory describes this as irreducible causal and informational structure [19]. Global-workspace theory describes it as system-wide availability [20]. Neural binding studies describe it as coordinated wholes [21, 22].

Why do we treat SU(3) as hard to break? In the thermodynamic, the low-temperature limit favors protected order and suppresses accessible disorder. The third law gives this limit its direction. Visible matter also shows strong stability at the proton level, since no proton decay has been observed. Free color charges are not observed either. When the strong sector is pulled apart, confinement forms new bound states instead of releasing isolated color. These facts make SU(3) a serious candidate for a remnant organizing structure. This motivates the use of the SU(3) confinement volume in Refs. [55, 56] to address the cosmological constant and the values of the fundamental constants.

### 5.2 Cessation as a Neural Ground Regime: Attenuating Electromagnetic Fluctuations

Cessation reports seem to define a boundary case quite clearly. The transition is marked by the absence of ordinary experience, including thoughts, perceptions, and the felt continuity of the self-model [39, 40]. This motivates the central question: what, if anything, is left, and what keeps it stable? This is where the SU(3) element of the argument enters. We treat it as a possible structure of what remains once the fluctuating degrees of self-organized experience are absent. If such an organizing layer exists, it would not be directly accessible. It would have to be inferred from transition regimes and stable composite observables. This is the same logic used in confinement physics, where the gluonic degrees of freedom of SU(3) are inferred through gauge-invariant composites rather than directly observed as isolated objects.

### 5.3 The Common Structure: Irreducible Residue After Expulsion of Fluctuations

The analogy is straightforward. In the cosmic realm, a superconducting-like vacuum repels destabilizing fluctuations. Third-law restrictions limit the remaining entropy, and the residual low-entropy order appears as an unbroken SU(3) framework built from confinement-scale vacuum elements. The neural side follows from the idea that the brain may also contain a superconducting-like link or channel. In deep cessation, organized electromagnetic and cognitive fluctuations become quiet. Experience moves toward a minimal-perturbation regime, and the residual integrated order may take the form of a compact, non-factorizing geometry. When reconstructed neural dynamics show the required generator structure, SU(3) becomes the natural candidate.

### 5.4 Related Theoretical Motivation: Observer-Dependent State Counting in Quantum Gravity

Related work in quantum gravity motivates the observer-dependent part of this framework. In de Sitter space, the Euclidean path integral for an isolated gravitational system can give imaginary contributions to the density of states. Maldacena showed that these terms disappear when the calculation includes a real observer with a worldline and a clock [57]. In that setting, the observer-inclusive calculation removes the imaginary contributions. This connects directly with the present proposal. Ref. [58] identifies the observer’s clock with an SU(3) system. In the vacuum-atom picture, the universe contains ~ 10^123^ SU(3) vacuum units whose collective organization fixes the observed dark-energy scale. The observer’s clock therefore tracks the same microscopic structure that organizes the vacuum. Thus, within this related theoretical lineage, de Sitter state counting, the vacuum-atom model, and the neural ground-regime analogy all point to the same principle: when fluctuating degrees of freedom are suppressed, the remaining order is organized by a stable, irreducible SU(3) structure.

## 6 SU(3) as the Groundless Ground: A Structured and Testable Framework

We now translate the two-fluid model into a symmetry model. The two-fluid picture separated ordinary dissipative activity from a coherence-supporting component. The Cartan-root decomposition of *su*(3) gives this split a precise algebraic form. The rank-two Cartan algebra Span{*T*_3_, *T*_8_} represents the slow coherence-balancing sector. These commuting directions preserve the integrated state and define the stable part of the dynamics. The six root generators represent transition directions. They change the reconstructed collective state and encode the flexible, content-changing part of the dynamics. This section therefore asks a narrow question: can neural data, after state-space reconstruction, reveal a two-plus-six generator structure compatible with *su*(3)? Raw EEG, MEG, ECoG, or impedance traces remain measured signals; the color-like variables below enter only as fitted collective coordinates. Section 7 then turns this symmetry model into a falsification test.

Geometrically, the claim is a connection claim rather than a gravitational claim: just as the Christoffel connection defines parallel transport on the tangent bundle in Riemannian geometry, a gauge potential 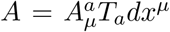 defines parallel transport on an internal SU(3) bundle, with curvature *F* = *dA* + *A* ∧ *A*. The present model uses this connection-geometric analogy for reconstructed neural state dynamics without implying literal spacetime gravity in the brain.

### 6.1 Model assumptions and reconstructed state space

Let *X*(*t*) denote the measured multichannel data stream, including EEG/MEG/ECoG, impedance, magnetic response, phase-locking structure, and other state-dependent markers. A reconstruction map Φ sends the measured signal to a latent collective trajectory,

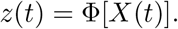

The data determine the dimension and geometry of this trajectory. The SU(3) hypothesis enters only if the recovered dynamics support an effective eight-generator closure compatible with *su*(3). Binding is then modeled as resistance to decomposition into independent sub-dynamics.

Let *C* denote the reconstructed collective state manifold when this closure holds. The variables below are inferred dynamical coordinates, not sensory channels or primitive contents. Phantom-limb phenomena, sensory substitution, and cross-modal recruitment after blindness show that perceptual organization can shift across neural substrates while retaining coherent structure [59–61]. Here they motivate the use of collective coordinates, without establishing the SU(3) claim.

### 6.2 From binding to a compact simple Lie-group model

If the reconstructed binding dynamics are compact, connected, non-factorizing, and eight-dimensional, the corresponding simple compact algebra is *su*(3). In the fundamental triplet representation, the candidate state manifold is

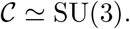

A single group element *U* ∈ SU(3) represents the integrated reconstructed state. The ground regime is the stationary limit

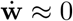

in the absence of external transients. The bi-invariant trace metric supplies the natural compact-group geometry. QCD confinement is used only as a structural guide: color degrees of freedom combine into non-factorizing observable composites; here distributed neural contributions are modeled as one reportable state.

### 6.3 Mathematical Realization: *su*(3), Group Elements, and the Metric

Let 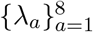 denote the (Hermitian) Gell–Mann matrices and define the standard *su*(3) generators by

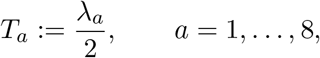

so that *iT*_*a*_ ∈ *su*(3) and

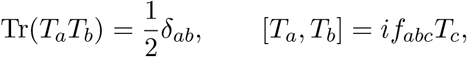

where *f*_*abc*_ are the usual SU(3) structure constants. We use the standard physics convention in which the Hermitian generators *T*_*a*_ = *λ*_*a*_/2 are identified with the *su*(3) basis; equivalently, the strictly mathematical anti-Hermitian basis is *iT*_*a*_.

A reconstructed collective state is represented by a group element *U* ∈ SU(3). The quantities *w*^*a*^ denote effective algebraic coordinates obtained after state-space reconstruction and basis fitting. They are mapped to real group coordinates *θ*^*a*^ = *θ*^*a*^(**w**), and the group element is then written as

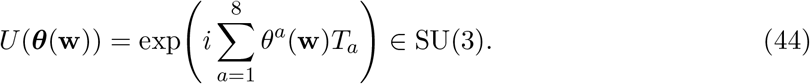

This is the compact representation of the global-binding condition.

The fundamental representation admits a temporal reading. Define

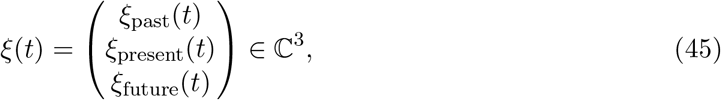

where *ξ*_past_ represents memory traces and past context structure, *ξ*_present_ represents present sensori-motor saliency and embodiment, and *ξ*_future_ represents future anticipation, predictions and intentions. The group element *U*(**w**) ∈ SU(3) transforms this triplet carrier as

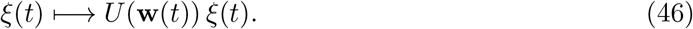

The triplet interpretation is a modeling choice placed on top of the algebraic test.

In QCD, color triplets *q* ∈ ℂ^3^ carry the fundamental representation. Here, the triplet *ξ*(*t*) ∈ ℂ^3^ carries temporal organization. The variables *w*^*a*^ are reconstructed algebraic coordinates. The empirical question is whether their effective transformations close approximately as *su*(3).

The triplet *ξ* = (*ξ*_past_, *ξ*_present_, *ξ*_future_)^*T*^ gives three temporal components. The Cartan sector gives two independent balance directions because SU(3) has rank two. Once unity fixes the full triplet, only two relative balances remain.

On the standard Gell-Mann basis,

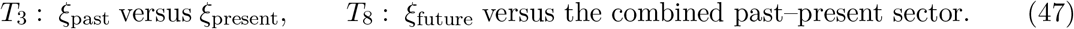

The data must first show a rank-2 commuting Cartan sector. The temporal reading then follows up to allowed basis changes.

The 6 real generators spanning the three root planes can thus specify the transition directions between the 3 components:

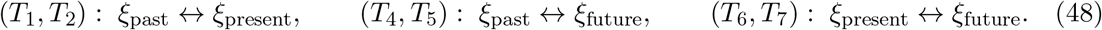

These root pairs define the candidate transition channels among memory, present experience, and anticipation.

Left-invariant representatives of the generators read

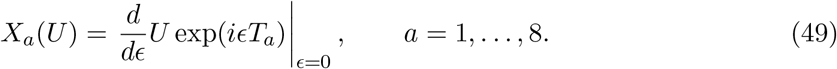

These vector fields give the transition directions of the reconstructed state manifold.

We define the inner product on *su*(3) by

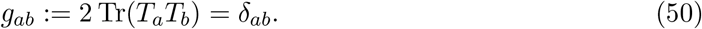

This defines the bi-invariant metric used below.

| Quantity | QCD role | Consciousness-model role |
| --- | --- | --- |
| Fundamental carrier | Color triplet $q = (q_r, q_g, q_b)^T \in \mathbb{C}^3$ | Effective temporal-conscious triplet $\xi = (\xi_{\text{past}}, \xi_{\text{present}}, \xi_{\text{future}})^T \in \mathbb{C}^3$. |
| Symmetry group | Local $\text{SU}(3)$ color gauge symmetry | Effective $\text{SU}(3)$ state-manifold symmetry acting on the temporal-conscious carrier. |
| Generators | $T_a = \lambda_a/2$ acting on color space | The same abstract generators represented as infinitesimal transformations mixing and rebalancing the triplet components. |
| Gauge fields (algebraic coordinates) | Gluon fields are algebra-valued gauge components | $w^a$ are reconstructed algebraic coordinates, not gluon fields or directly measured observables. |
| Cartan sector | Two diagonal generators associated with color-space weights | Two independent coherence-preserving balance relations among the three triplet components. |
| Root sector | Six off-diagonal generators mixing color components | Six transition-generating directions: memory–present updating, memory–future projection and present–future anticipation/action-readiness. |
| Empirical test | Hadronic observables and QCD scattering/structure data | Reconstruction of neural dynamics and testing of the temporal-triplet Cartan–root algebraic signature. |

### 6.4 Cartan–root realization of the two-fluid split within the SU(3) model

The two-fluid picture separates coherence-supporting and transition-generating dynamics. The Cartan–root decomposition gives this split an algebraic form: Cartan directions preserve balance; root directions generate transitions.

The superconducting analogy motivates the split. In the coherent regime, the charged condensate acquires ⟨Ψ⟩ ≠ 0, the electromagnetic response becomes Higgsed, and the Meissner effect screens long-range magnetic fields. The neural model uses SU(3) as a compact non-Abelian organizer of the reconstructed state space.

Let 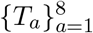, with *T*_*a*_ = *λ*_*a*_/2, denote the Gell–Mann basis. The Lie algebra *su*(3) admits the standard decomposition

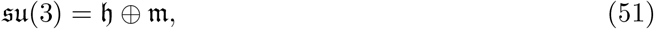

where the Cartan subalgebra is

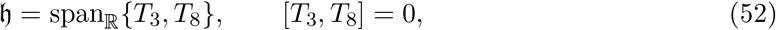

and the root-space complement is

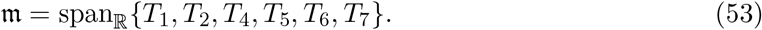

With respect to the bi-invariant inner product ⟨*X, Y* ⟩ = 2 Tr(*XY*), this is an orthogonal direct sum. Its basic bracket structure is

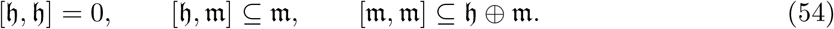

Thus *h* carries commuting directions, while *m* carries non-Abelian transitions.

For the state element *U*(***θ***(**w**)) of Eq. (44), define the Hermitian algebra-coordinate velocity

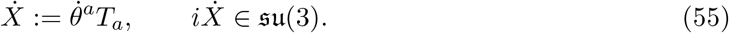

Let *P*_*h*_ and *P*_*m*_ denote the orthogonal projectors onto *h* and *m*. Then

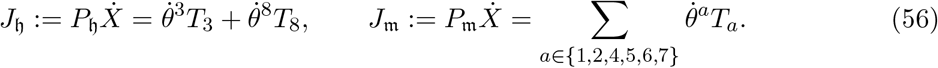

Here *J*_*h*_ gives the coherence-supporting sector, and *J*_*m*_ gives the transition-generating sector. The sectional curvature formula for a bi-invariant metric,

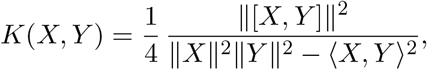

vanishes for *X, Y* ∈ *h*, so the Cartan sector moves on a flat maximal torus *T* ^2^ ⊂ SU(3). The root sector carries the non-Abelian curvature of the group.

The rank gives the balance directions, while dim *G* − rank *G* gives the transition directions. For several compact connected simple Lie groups one obtains

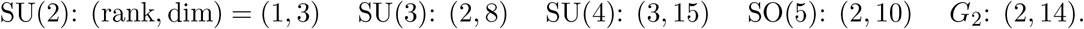

SU(3) is the smallest option in this list with rank two and an eight-generator non-factorizing algebra. Define the sector norms

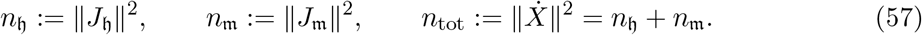

In the unforced bi-invariant geometric limit, *n*_tot_ is conserved along geodesic motion. Within the model, *n*_*h*_*/n*_tot_ measures the coherent fraction, while *n*_*m*_*/n*_tot_ measures the transition-generating fraction.

The empirical content is the algebraic signature: a rank-2 approximately commuting sector and a structured six-dimensional transition sector in reconstructed neural dynamics. If independent datasets lack this Cartan–root structure, the interpretation loses support. The two Cartan directions may describe two balance modes of conscious time, such as retention and anticipation [62]; formally, only the rank-2 commuting sector is required.

### 6.5 Intrinsic Geometry and a Model Field Equation for Internal Dynamics

The bi-invariant metric equips SU(3) with a symmetric intrinsic geometry. Standard Lie-group geometry yields an Einstein-manifold relation of the form Ric_*ab*_ ∝ *g*_*ab*_ for compact simple groups equipped with their bi-invariant metric. In the normalization adopted here, we write:

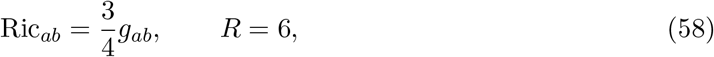

which fixes the corresponding Einstein tensor.

To connect geometry to coarse-grained dynamics, introduce a model current in the reconstructed algebraic coordinates:

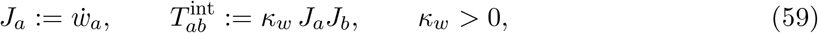

and postulate the internal equation

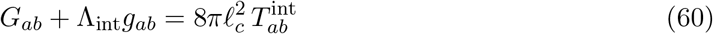

as a minimal geometric closure. This equation is used only as an internal connection-geometric closure on the reconstructed state manifold, not as a claim of literal gravitational dynamics in brain tissue. Here *ℓ*_*c*_ is an assumed mesoscopic length scale of the relevant integrative substrate (taken as *ℓ*_*c*_ ≈ 10^−5^ m to match a mesoscopic cortical scale [63, 64]), and Λ_int_ sets the vacuum curvature scale.

Using (58), the Einstein tensor evaluates to

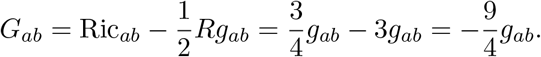

In the stationary regime 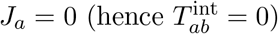, equation (60) yields the vacuum condition

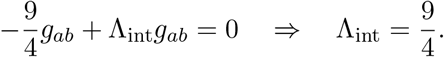

Thus the stationary model state 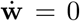 corresponds to the vacuum solution of (60) with Λ_int_ = 9/4. Cessation-like reports motivate this limiting case [39], but the mapping remains a model assumption.

### 6.6 Multi-Timescale Neural Dynamics on a Reconstructed SU(3) Arena

The SU(3) manifold supplies the arena for the reconstructed collective state **w**(*t*). Additional neural variables steer **w**(*t*) across the manifold. On fast neural timescales of order 10 ms, the high-dimensional neural state **x**(*t*) ∈ ℝ^*N*^ evolves according to 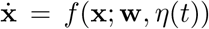, where *η*(*t*) represents transient inputs and fluctuations. On slower cognitive timescales, from seconds to minutes, the reconstructed algebraic state evolves as 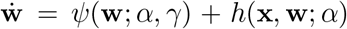. On still longer timescales, meta-parameters drift according to 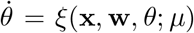. These variables drive the trajectory on the proposed group-theoretic arena; they do not fix the decomposition of experience.

### 6.7 Extreme Regimes and Altered States as Geometric Signatures

Equation (60) organizes altered regimes by the magnitude and anisotropy of the current 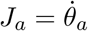, with *θ*^*a*^ = *θ*^*a*^(**w**). Ordinary wakefulness has sustained modulation, *J*_*a*_ ≠ 0. Absorption or Samādhi-like regimes are represented by reduced or more structured currents [65]. Cessation is treated as a non-phenomenological boundary case, modeled by suppression of reportable variation and an approach toward *J*_*a*_ → 0 [39, 40]. Anesthesia corresponds to altered EEG signatures of loss and recovery of consciousness [66]. Psychedelic states correspond to broader state-space exploration and altered integration signatures [67, 68].

### 6.8 Mathematical status of the model

The role of SU(3) is conditional. If reconstructed neural dynamics support a compact, non-factorizing, eight-generator simple algebra, Cartan classification selects *su*(3). The stationary regime 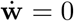 defines the model ground state, and (60) represents it as a vacuum solution with Λ_int_ = 9/4. The empirical assumptions are not mathematically forced. They define a specific algebraic target that neural data can support, weaken, or falsify.

## 7 Operational test of the rank-2 Cartan signature

This section turns the model into a deliberately hard empirical test. The prediction may fail in real neural data; that is the point. A resistance drop, a coherence increase, or a two-mode neural manifold would not by itself support SU(3). The model requires three linked findings: an eight-dimensional reconstructed effective algebra, a rank-2 approximately commuting slow sector, and a six-direction transition sector whose commutators resemble *su*(3).

The physical layer and the symmetry layer must be separated. Resistance, impedance, non-linear transport, magnetic response, phase stiffness, and heat capacity test whether a collective low-dissipation component exists. They do not test the SU(3) algebra. The algebraic test begins only after state-space reconstruction.

Let the measured multichannel data be written schematically as

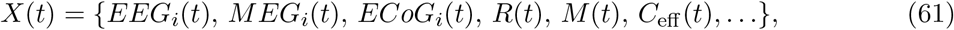

The measured signal is *X*(*t*). The extra material readings are *R*(*t*), *M*(*t*), and *C*_eff_ (*t*). From the signal we build a path in state space,

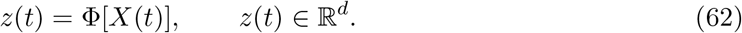

The map Φ can be built with standard tools for brain-state reconstruction [68–70]. The dimension *d* is chosen from the data before the symmetry test is applied. The SU(3) reading needs *d* = 8. A stable preference for another value of *d* weakens that reading.

Suitable inputs include neural time series, phase-locking metrics, microstate occupation probabilities, impedance or resistivity measurements, magnetic response, phase-stiffness proxies, heat capacity, and transition rates between states. These should be compared across wakefulness, absorption-like states, fragmented states, anesthesia, and unconscious controls. The decisive tests are rank two, approximate commutation, and a six-direction transition sector with empirical commutators close to the *su*(3) structure constants.

A convenient explicit implementation is the following. When an *su*(3)-compatible reconstruction is supported, form the data matrices

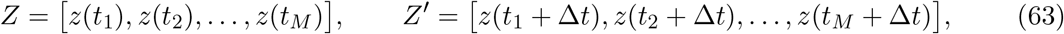

and estimate a linear propagator on the reconstructed space by

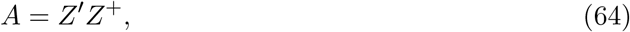

where *Z*^+^ is the Moore–Penrose pseudoinverse. Dynamic mode decomposition gives

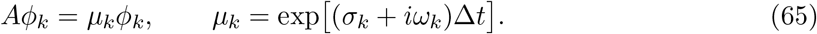

Here *σ*_*k*_ is a damping or growth rate and *ω*_*k*_ is an oscillation frequency. Sorting the modes by increasing |*σ*_*k*_| identifies the slow sector. The observed slow rank is then

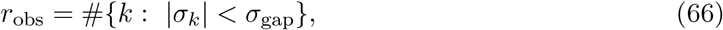

where *σ*_gap_ is fixed by a pre-registered spectral-gap criterion. If no stable slow sector exists, the SU(3) interpretation fails at this stage.

If the reconstruction passes this first filter, the trajectory is tested against an SU(3)-structured representation,

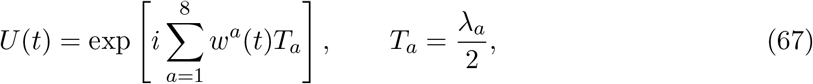

where *λ*_*a*_ are the Gell-Mann matrices. The instantaneous group-space velocity is

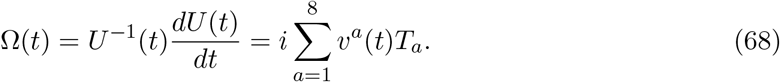

The Cartan and root components are then

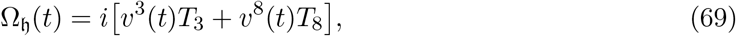

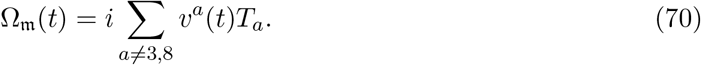

The first measurable statistic for Cartan is the coherent Cartan fraction

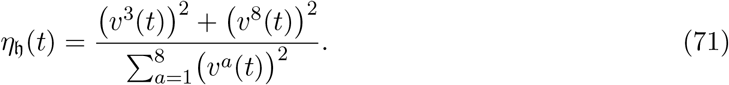

This statistic is only a screen. It becomes meaningful after basis rotation, slow-sector stability checks, and comparison with controls.

The statistic *η*_*h*_ measures the fraction of instantaneous motion carried by the rank-two slow sector. Alone, it gives a weak test. The SU(3) test also requires approximate commutation and the expected structure-constant pattern. Thus the rank-two sector must dominate coherent dynamics and close algebraically. For that last property, one wants to have

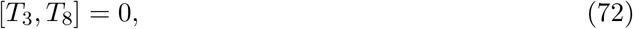

so the two modes *V*_1_, *V*_2_ reconstructed should satisfy

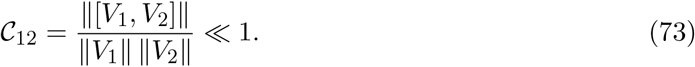

Operationally, applying the two reconstructed flows in opposite orders should return nearly the same state.

The stronger test uses the full algebra. We reconstruct effective generators 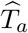 on the recovered state space. We then compare their empirical commutators with the *su*(3) structure constants. Define

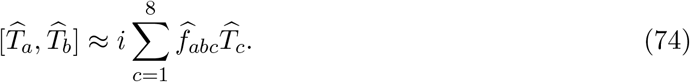

Then an SU(3)-specific mismatch statistic may be written as

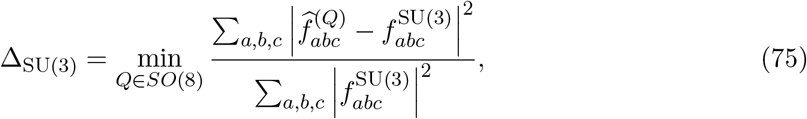

Here *Q* rotates the reconstructed basis. The *SO*(8) fit should be repeated from many starting points and benchmarked against simulated, shuffled, and competing-algebra controls.

The method should first be tested on simulated SU(3) dynamics with EEG- or MEG-level noise. Rank 2 alone is too broad. The commutator pattern gives the real SU(3) test. The expected reconstruction can be summarized schematically in Fig. 8.

**Figure 8:**
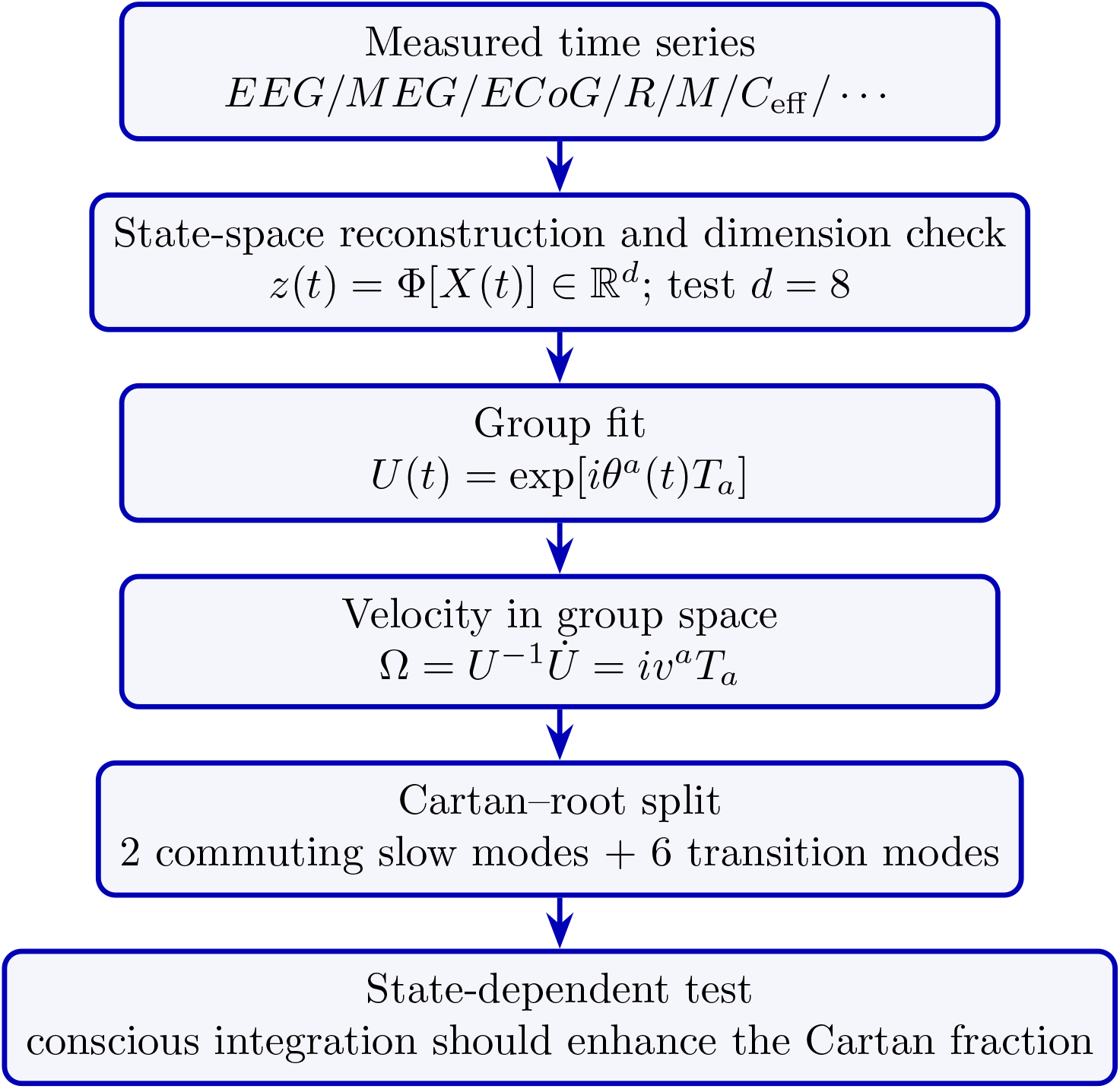
Operational reconstruction pipeline for the proposed SU(3) test. Physical transport and magnetic observables enter the measured data vector, but the symmetry test is performed only after reconstructing the collective state-space dynamics and testing the Cartan–root split. The symmetry test in the final box is independent of the physical-layer measurements in the top box and uses only the reconstructed coordinates.

The corresponding phase-space distinction is illustrated in Fig. 9, which separates the proposed rank-2 Cartan signature from rank-1, generic two-mode, and noisy alternatives.

**Figure 9:**
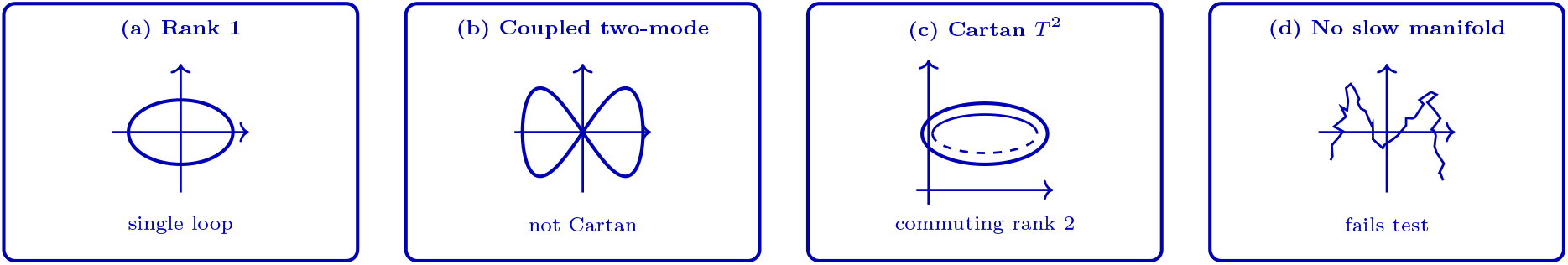
Operational phase-space distinction. A single resistance or pendulum-like loop gives a rank-1 picture; a generic two-mode system need not be Cartan; the SU(3) prediction is a rank-2 approximately commuting slow sector, represented schematically by a flat two-torus *T* ^2^; and a noisy state with no stable slow manifold fails the symmetry test.

Current neuroscience supplies relevant comparison classes: neural manifolds [69], EEG microstates [22], brain thermodynamics [3], absorption, and anesthesia [46, 66]. These sources motivate the reconstruction strategy, not the SU(3) result.

The transition-vector histogram gives a useful visual tool. After reconstruction, define small increments:

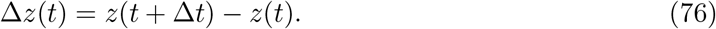

We project these increments onto the transition directions and remove the two Cartan directions, then record the resulting angles in a histogram. The expected outcome of the SU(3) model is a six-direction structure in the transition histogram. This pattern is weakened by a flat angular distribution. While simple to inspect, the transition histogram is insufficient by itself. A stronger test uses the commutators and structure constants. A suitable decision rule is summarized as follows:

| Observation | Interpretation |
| --- | --- |
| No slow manifold | The SU(3) interpretation is not supported. |
| If $r_{\text{obs}} = 1$ or $r_{\text{obs}} \geq 3$ | The recovered dynamics does not have the hypothesized 2-dimensional Cartan sector. |
| If $r_{\text{obs}} = 2$, but the Cartan-pair fidelity is poor | There is a linear two-state system present, but no true Cartan pair. |
| If $r_{\text{obs}} = 2$, with good fidelity and the right structure of the 6-dimensional transition sector | The Cartan-root interpretation is strengthened. |
| If the same signatures are seen in shuffled or controlled signals | The consciousness-specific aspect of the interpretation is weakened. |
| If the signal improves as subjects transition from unconscious to integrated conscious states and disappears under anesthesia | The SU(3) interpretation strengthens. |

The slow-manifold dimension gives only the starting point. The structure constants carry the actual SU(3) test. The mismatch statistic should therefore be evaluated against simulations, shuffled controls, and competing symmetry models under realistic EEG or MEG noise. A null result would not damage the general study of neural manifolds; it would specifically weigh against this SU(3) interpretation.

The resistance phase-space plot probes the physical layer. The SU(3) interpretation requires reconstructed neural dynamics. Absence of the rank-2 Cartan signature and six-direction transition structure would weigh against the model. Observation of the full pattern during integrated conscious states, and its loss under anesthesia or unconscious controls, would strengthen it.

## 8 Conclusion

One of the core tasks of the brain is managing segregation (e.g., the visual and tactile input of a cup as a separate object with particular characteristics) and integration (i.e., the coherent, unified state of consciousness that includes the body, mind, and cup). The other fundamental challenge is orchestrating this complex model of the self, body, and the world on a highly limited energy budget, approximately 20 watts. Hence, the brain must somehow maintain and balance a rapid, coherent, integrated state with a more coarse-grained and abstract capacity for representation and conceptualization; and it must do so extremely rapidly and efficiently.

To address these open challenges and others, we developed a unified theoretical proposal linking SU(3) symmetry with superconductivity-inspired coherence under the thermodynamic protection of low-entropy regimes. We propose that unified conscious organization should be treated as a reconstructed collective dynamical state, and the SU(3) structure is proposed as a candidate algebraic geometry for its non-factorizing integration.

Moreover, superconductivity is proposed to supply a candidate mechanism by which efficient, large-scale integration might be supported under biological conditions. Specifically, we draw an analogy from the two-fluid paradigm and modeled neural dynamics as the interaction of a dissipative component and a coherence-supporting component, where different states of consciousness and cognition correspond to changes in their relative contribution. In other words, consciousness may be a unified, dynamically evolving pattern of brain activity, integrated through superconductivity-like coherence, whose internal organization can potentially be described using SU(3) symmetry.

The framework is also falsifiable. The Lie algebra *su*(3) contains a rank-two commuting sector and six root directions, which we interpret as relatively stable, coherence-preserving dimensions and transition-generating dimensions, respectively. Effective generators reconstructed from neural trajectories should exhibit a robust two-plus-six organization across changes in conscious state and should explain the data better than competing dynamical models. In the future, we should also see further independent evidence of the signatures of superconductivity in microtubules.

## Acknowledgements

We gratefully acknowledge Dr. Miranda DeWitte for providing very helpful comments and references on the interconnected role of blood pressure, theta waves, electrical conductivity, temperature and anesthetic agents in the context of consciousness. The authors also gratefully acknowledge support from Texas Space LLC (texasspacellc.com) and Juan Feria. The content of this paper is being used to develop associated patents by Texas Space.

## Footnotes

1 This Lagrangian density is closely related to *φ*^4^ theory and the Abelian Higgs model, which have deep implications in particle physics, specifically, particles acquire mass due to spontaneous symmetry breaking (e.g., the Higgs mechanism). [28, 29]

2 Note that Wilczek [30] refers to this as “gauged broken symmetry” versus “broken gauge symmetry.” The reason is because local gauge invariance cannot be broken. Rather, in a model of broken global symmetry, the mass term arises.

3 Note that we could also use the Hilbert (metric) stress tensor: 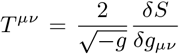. In flat space-time, this becomes 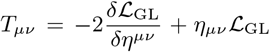. Using the Lagrangian in (1) leads to the same energy-momentum-stress tensor obtained in (18).

4 For other discussions of superconductivity as a condensed-matter analog of *U* (1) gauge symmetry breaking and the Higgs mechanism, see section 21.6 of Weinberg’s text [34], and sections 8.3 and 8.4 of Ryder’s text [28], and p. 41 of [35]. For an insightful alternative perspective on this point, see [36].

